# Exploring cell surface-nanopillar interactions with 3D super-resolution microscopy

**DOI:** 10.1101/2021.06.21.449280

**Authors:** Anish R. Roy, Wei Zhang, Zeinab Jahed, Ching-Ting Tsai, Bianxiao Cui, W.E. Moerner

## Abstract

Plasma membrane topography has been shown to strongly influence the behavior of many cellular processes such as clathrin-mediated endocytosis, actin rearrangements, and others. Recent studies have used 3D nanostructures such as nanopillars to imprint well-defined membrane curvatures (the “nano-bio interface”). In these studies, proteins and their interactions were probed by 2D fluorescence microscopy. However, the low resolution and limited axial detail of such methods are not optimal to determine the relative spatial position and distribution of proteins along a 100 nm-diameter object, which is below the optical diffraction limit. Here, we introduce a general method to explore the nanoscale distribution of proteins at the nano-bio interface with 10-20 nm precision using 3D single-molecule super-resolution (SR) localization microscopy. This is achieved by combining a silicone oil immersion objective and 3D double-helix point-spread function microscopy. We carefully optimize the objective to minimize spherical aberrations between quartz nanopillars and the cell. To validate the 3D SR method, we imaged the 3D shape of surface-labeled nanopillars and compared the results with electron microscopy measurements. Turning to transmembrane-anchored labels in cells, the high quality 3D SR reconstructions reveal the membrane tightly wrapping around the nanopillars. Interestingly, the cytoplasmic protein AP-2 involved in clathrin-mediated endocytosis accumulates along the nanopillar above a specific threshold of 1/R membrane curvature. Finally, we observe that AP-2 and actin preferentially accumulate at positive Gaussian curvature near the pillar caps. Our results establish a general method to investigate the nanoscale distribution of proteins at the nano-bio interface using 3D SR microscopy.

## Introduction

The cell-to-material interface is often a key determinant of successful applications for tissue engineering and biomedical implants. Material properties, including chemical functionalization, surface topography, and bulk stiffness, collectively set instructive signals for cell behavior^1, 2^ Nanoscale surface topography is particularly interesting as it is widely tunable and has been shown to significantly affect cellular responses. In the last decades, nanofabrication emerged as a powerful tool to precisely engineer nanostructures, i.e. the nano-bio interface, to control cell behavior. For example, nanopillars have been shown to reduce focal adhesions and membrane tension^3, 4^; nanogratings and nanofibers induce cell alignment and neural development^5, 6^; and nanopores accelerate stem cell differentiation^7, 8^. Nanopillars made of different materials were also developed into electrical and optical sensors for measuring live cell activities^9, 10^. Due to its biological significance, there is great interest to visualize the nano-bio interface, especially with respect to the membrane shape around nanotopography and intracellular proteins at the interface. By varying the diameter of nanopillars that changes the shape of the membrane at the interface, recent studies show that nanopillars locally activate clathrin-mediated endocytosis and the polymerization of actin fibers in a curvature dependent manner^11, 12^. However, visualizing the membrane shape at the interface and quantifying the curvature values are technically challenging, especially in three dimensions (3D).

Ultrastructural analysis by electron microscopy remains a powerful approach for visualizing the nano-bio interface. Transmission Electron Microscopy (TEM) has been used to visualize the membrane shape around nanopillars and to measure the gap distance at the interface. Focused-ion-beam milling and scanning electron microscopy imaging (FIB-SEM) can be more advantageous by allowing selective opening and imaging of the interface at desired locations^13^.

However, as is well-known, electron microscopy gives a snapshot of the physical effects with very high spatial resolution and general cellular context, but without specific identification of non-discernable proteins, while fluorescence microscopy gives molecular specificity without going beyond the optical diffraction limit (DL) of 250-300 nm. Recent work in correlated single-molecule localization and cryogenic electron tomography has shown important progress in combining these two modalities^14^. Previous studies have used fluorescence 2D DL experiments to study the behavior of specific proteins at the nano-bio interface. However, analyzing the nanoscale distribution of the proteins around and along the entire nanopillar in 3D and at 10-20 nm resolution is not possible from to the low resolution 2D DL images. While some experiments have probed the nano-bio interface at higher resolutions than the DL, more information would be obtained about the nanoscale distribution along the pillars with 10-20 nm resolution in 3D^15, 16^. To address this optically, we describe a method to use super-resolution fluorescence microscopy (SR) in three dimensions (3D) to precisely explore the positions of probe molecules down to ~10 nm in cells interacting with fabricated quartz nanopillars. Our approach required optical optimization and the use of control imaging tests to validate the procedures. We show that membrane-anchored labels as well as proteins that preferentially accumulate on curved membranes such as AP-2 and actin interact with the curvature constraints of the nanopillar substrate in different ways. This work shows the utility of the 3D SR approach and should stimulate further use of the method to quantitatively characterize the nano-bio interface.

## Results and discussion

### Nanopillar imaging strategy to mitigate spherical aberrations

Recent studies used a standard oil immersion objective (OIO) in 2D DL fluorescence imaging experiments to image the behavior of cytoplasmic proteins such as AP-2 and actin on quartz nanopillars^11, 12^. Moving from 2D DL to 3D SR imaging on the quartz nanopillar substrate is not straightforward. The standard OIO is optimized for imaging through glass rather than quartz substrates because the refractive index of the immersion oil matches the glass refractive index. In general, any refractive index mismatch between the cover slip and the sample will lead to spherical aberrations that deteriorate image quality. Specifically, the problem at hand can be affected by the refractive index mismatches between the cell (n~1.40), the quartz substrate and nanopillars (n=1.45), and the immersion oil (n=1.52). Due to the limited spatial resolution of 2D DL, the effect on image quality or the resolution is not very significant or observable in most images of the interface. However, the 3D localization precision and localization accuracy will degrade rapidly in the presence of spherical aberrations and will deteriorate the final SR reconstruction image quality^17–23^.

To mitigate spherical aberrations arising from imaging cells on quartz nanopillar substrates with 3D SR, we repurposed a silicone OIO with a correction collar in order to more carefully match the refractive index mismatches. Similar to water immersion (WI) objectives that index match to aqueous media with a carefully chosen glass substrate thickness, the silicone OIOs strive to use the approximate match between the silicone oil (n=1.40) and the index in the cell cytoplasm. As a result of the objective design, the optical path mismatch induced by a glass coverslip can be reduced by carefully adjusting the correction collar. The silicone OIO has been employed successfully in various microscopy methods^24, 25^. Here we show that the adjustable silicone OIO can also work well with the cells-on-quartz nanopillar 3D SR imaging problem.

Figure 1A depicts the advantages of using a silicone OIO compared to the standard OIO for a simple case where a cell is imaged on a flat quartz coverslip. With the standard OIO, light refracts between the cell cytoplasm and the coverslip and between the coverslip and oil. Ultimately, the wavefront distortion from the refraction results in the spherical aberrations^26^ that degrade localization precision and accuracy. However, with the calibrated silicone oil objective, the light experiences minimal refraction at the interfaces, and there is less wavefront distortion, less spherical aberration, and higher image quality. The only issue here is whether the silicone OIO can be corrected sufficiently for our quartz substrate, given the objective design assumes glass coverslips.

**Figure 1:**
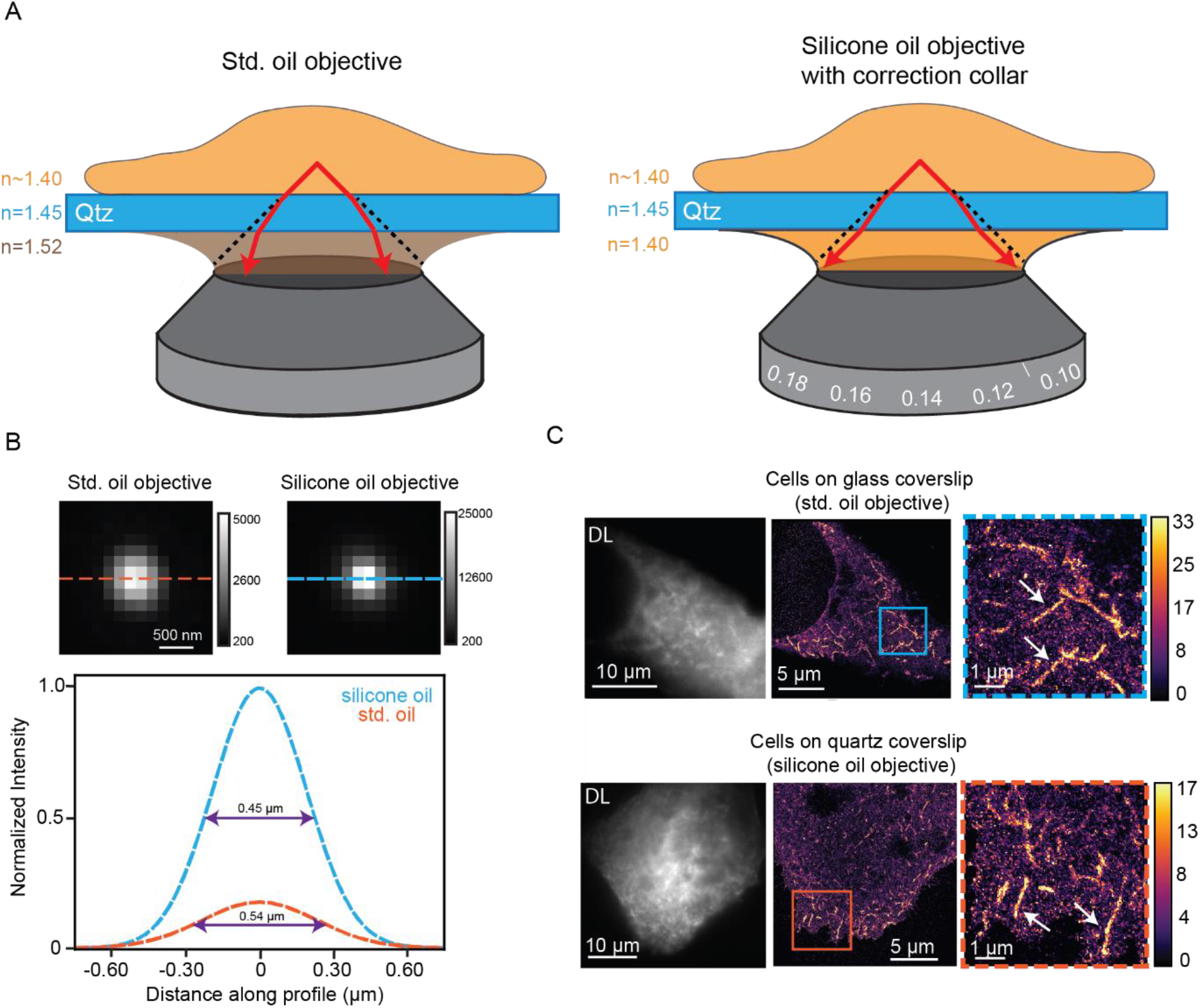
Comparison of silicone OIO standard OIO (std.). (A) Cartoon schematic depicting how light rays refract using the std. OIO (left) and the silicone OIO on quartz (Qtz) substrates. (B). Images of 200 nm beads immobilized in 5% agarose on quartz coverslips imaged with both objectives (top). Calibration bar depicts ADC counts on the EMCCD camera. Cross sections (orange and blue lines) are fit to a Gaussian (bottom). The full width half maximum (purple double arrow) is clearly smaller for the case with the silicone OIO (C) DL (left) and 2D SR reconstructions (middle and right) of FBP-17 labeled cells imaged with the std. OIO and silicone OIOs grown on glass and quartz substrates respectively. Calibration bar depicts number of localizations in each bin of the histogram reconstruction (bin size 32 nm). Orange and blue boxes are magnified images of the 2D SR reconstructions. White arrows show examples of tubule invaginations.

The correction collar of the silicone OIO is first carefully tested on a 200 micron thick quartz coverslip, the thickness used to fabricate nanopillars in later experiments. To do this, we imaged 200 nm poly(styrene) fluorescent beads immobilized in 5% agarose on 200 micron thick quartz coverslips at various collar adjustments, and the extent of spherical aberrations at each adjustment was assessed by quantifying the peak intensity of the bead image spots. Spherical aberrations will reduce peak intensity because the photon distribution is spread over a larger spatial scale^27, 28^. The peak intensity of the observed spot is a common metric used to assess the extent of spherical aberrations present in the image and has been used in approaches such as adaptive optics^29, 30^. We set the correction collar to the adjustment with the maximum peak intensity (Methods and Figure S1A). This adjustment was used for all imaging experiments involving the 200 micron thick quartz substrate.

Figure 1B depicts representative beads on 200 micron thick quartz substrates imaged with both the standard and silicone OIOs. The bead imaged with the silicone OIO has a more tightly focused photon distribution and thus higher peak intensity compared to the bead imaged with the standard OIO. Horizontal cross sections fit to a Gaussian (plots below bead images) also show that the full width half maximum is smaller for the bead imaged with the silicone OIO. We conclude that fewer spherical aberrations are present leading to expected superior performance with the silicone OIO compared to the standard oil OIO. Quantifying the spherical aberrations over many beads imaged with both objectives further confirms the approach (Figure S1B).

Imaging on quartz substrates with a silicone OIO can then be extended to cellular SR imaging. We first chose to verify that the SR image quality from cells adhering to quartz substrates using the silicone OIO are comparable to the quality from standard imaging approaches: imaging cells on glass substrates with the standard OIO. We overexpressed the protein Formin-binding protein 17 (FBP17) in U2OS cells and grew cells on glass and quartz substrates. FBP17 is a Bin/amphiphysin/Rvs (BAR) domain protein, a class of proteins that are banana-shaped, bind preferentially to regions of membrane curvature, and can also induce membrane curvature^31–33^. When overexpressed in cells, FBP17 forms tubule invaginations that are not resolvable using DL imaging^34^. The ability to resolve the tubule invaginations will be used to compare the image quality of both imaging modalities.

FBP17 was expressed with a green fluorescent protein (GFP) domain fusion and was labeled with GFP nanobodies covalently labeled with Alexa Fluor 647 (AF647, Figure S1C for controls). 2D Stochastic Optical Resolution Microscopy (STORM^35^, dSTORM^36^) SR data was acquired by imaging fixed cells close to the coverslip with high laser intensity in a blinking buffer. We also call this type of SR microscopy SMACM for single-molecule active control microscopy^37^ as a more general term for any mechanism that forces the concentration of emitting molecules in single frames down to a sparse level allowing for single-molecule localizations and subsequent SR reconstruction. The data was then processed to yield 2D SR reconstructions. Figure 1C (first column) depicts the DL images of labeled FBP17 cells in both objective/substrate combinations. Due to the diffraction limit, the features of the invaginations are not easily apparent. The second and third columns show the 2D SR reconstructions of the cells first on a large scale and then as magnified images, respectively. The features of the tubule invaginations in the SR reconstructions are much more clearly observed and, qualitatively, the invaginations appear similar in both situations. In addition, the median localization precision is 10 nm (Figure S1D for distributions) for both cases, and quantification of the diameters of the invaginations (Figure S1E) is similar for both configurations and is in close agreement to literature^38, 39^. Thus, as the image quality is both qualitatively and quantitatively similar for both imaging configurations, 2D SR imaging of cells on quartz substrates with the silicone immersion objectives retains the image quality found using standard SR imaging approaches.

### 3D SR microscopy of surface-labeled nanopillars using a double-helix PSF microscope

Next, the imaging configuration combing a quartz substrate and silicone OIO must be adapted for 3D SR imaging of the nano-bio interface. Even with standard open-aperture widefield microscopy, the shape of the detected single-molecule spots, regarded here as the point spread function (PSF), changes as a function of defocus. However, extracting 3D position from the shape changes is challenging. The shape of the standard PSF is symmetric above and below focus, resulting in potential redundancy of the Z position. In addition, the shape quickly blurs 400 nm away from focus in both directions, and determining Z requires high signal-to-noise^40^. As cells and the nanopillars may extend in the axial direction several microns, the relatively short imaging range of ~800 nm is not desirable.

To circumvent these issues, we used PSF engineering approaches to more optimally extract Z-position, as described in many previous studies^41–43^. In PSF engineering, we insert a simple transmissive phase mask in the Fourier plane (conjugate to the back focal plane) of the microscope. As the Fourier plane is usually found close to the back of the objective in many microscopes, it can be challenging to access. Thus, we used simple 4f emission processing optics outside the microscope to relay the collected fluorescence light from the usual image plane to a new image plane four focal lengths away. Now there is easy access to the Fourier plane as illustrated in Fig. 2A. The phase mask imparts a phase delay in the collected fluorescence light that modulates the shape of the PSF after the light is focused on the camera detector. Here we chose to insert a double-helix phase mask^41^ in the Fourier plane of our microscope (phase pattern Fig 2A inset), which has been used in previous 3D SR imaging experiments with cells^21, 44, 45^. The modified PSF now has two lobes and is termed the double-helix PSF (DHPSF)^41^. The axial range of this DHPSF design is 2 microns, the shape is asymmetric above and below focus, and it changes rapidly with Z, circumventing the issues of the standard PSF for 3D imaging and facilitating precise Z position estimation. The XY position is extracted by fitting the DHPSF spot on the camera (see Fig 2D for examples) to a double-Gaussian function and finding the midpoint of the fit. As the angle between the lobes rotates as a function of the Z position of the emitter, a carefully calibrated curve measured prior to data acquisition connects the lobe angle to Z position. We note that no scanning of the objective or the stage occurs during data acquisition. The focus is simply set to one position, and we acquire DHPSF images of all emitters over the entire axial range of the DHPSF and extract the Z position in post processing. This localization estimation procedure has been shown to provide localization errors independent of Z, as opposed to other approaches to 3D such as astigmatism^43^.

**Figure 2:**
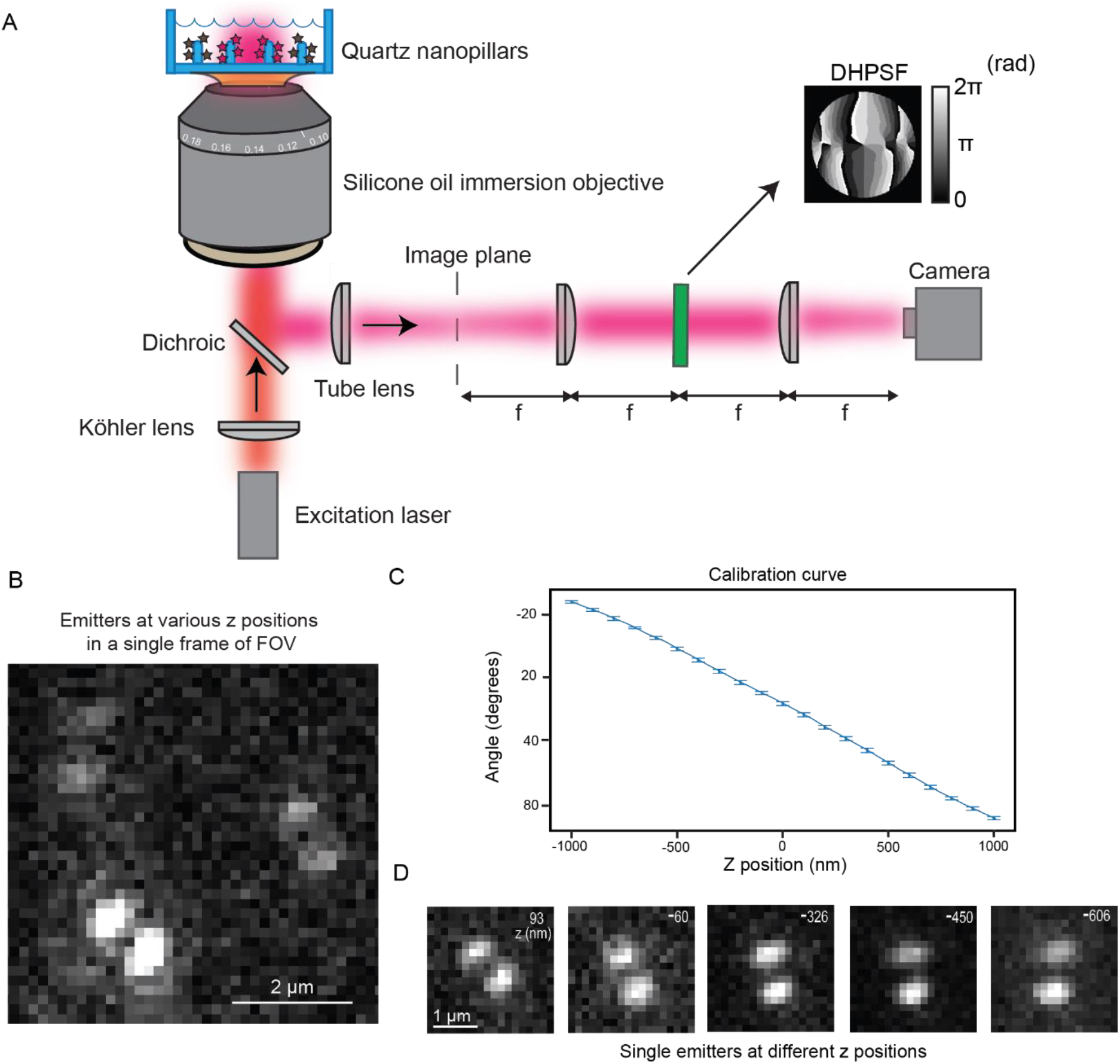
Schematic of microscope and representative images of single molecules imaged with the DHPSF. (A) Cartoon depiction of our DHPSF microscope. Excitation light entering the objective in the epifluorescence configuration is used to image surface labeled nanopillars. The emission is relayed through 4f emission optics onto the camera detector. The DHPSF mask (mask shown by arrow) is inserted in the Fourier plane for 3D imaging. Calibration bar in units of radians. (B) Representative field of view of experimental data showing three emitters. As the lobe angles for all three emitters is varied, the Z positions are different. (C) This plot depicts the calibration curve that correlates the lobe angle to the Z position. (D) Selected images reveal experimental DHPSFs extracted from a SMACM dataset of the surface-labeled nanopillars. The emitters were fit to extract the Z position (Z position shown in top right of image).

Using the silicone OIO and a DHPSF microscope, in principle we can now image proteins in 3D near the quartz nanopillars. However, a certain degree of spherical aberration may still be present and may potentially be Z-dependent. For instance, the light closer to the top of nanopillars experiences more refraction than the light near the bottom of the pillars. Therefore, we chose a model system to first benchmark the image quality of our 3D SR reconstructions. We will compare the dimensions and shapes extracted from 3D SR reconstructions of covalently surface-labeled nanopillars and from scanning electron microscopy (SEM) images of the nanopillars. If the image quality is high, the shapes and dimensions of the 3D SR reconstructions should be similar to the shapes and dimensions from the SEM images.

Fabrication of the nanopillars is described in the next paragraph, so we continue with 3D SR DHPSF imaging of covalently labeled quartz nanopillars here. To optically image these pillars, we first surface-labeled them by modifying the surface with free amine groups and then covalently attaching AF647 fluorophores using NHS chemistry with low nonspecific binding (Methods and Figure S2C for controls). We then collected many frames containing emitters bound to many pillars and within the axial range of the DHPSF. Figure 2B is a representative small field of view (FOV) in a frame of our acquired data where we see three emitters with various lobe angles and thus, different Z-positions. Figure 2C (top) depicts the calibration curve that connects the lobe angle to Z position of a poly(styrene) fluorescent bead on the surface; this was acquired by moving to known Z positions using a precise motorized piezo stage prior to SMACM data acquisition. Using this curve and double-Gaussian fits, we extract the 3D position of the emitter along the nanopillars for all the blinking single molecules from the camera image stack. Figure 2D showcases selected emitters at different Z positions. As the heights of the pillars are 884±72 nm (mean ± standard deviation), the entire axial range of the DHPSF is not fully utilized here. The total axial range of the emitters in Figure 2D covers the entire height of the pillar selected.

The other crucial aspect of our experiments is the fabrication of the quartz nanopillars, which follows previous work with some modifications. One modification utilized nanopillars fabricated with photolithography and chemical etching methods rather than electron beam lithography, in order to easily add a design containing a reference marker in an array pattern (Fig. 3A, described in Methods and Figure S2A). The reference marker allowed specific correlation between a pillar in the 3D SR reconstructions and the same pillar in the SEM image. To reduce the diameter of nanopillars to below the optical diffraction limit, we employed wet etching after the dry etching step. We used buffer oxide etchant for quartz substrate, which provides a generally isotropic process to remove material from the substrate and shrink material to a given dimension. This fabrication process resulted in tapering elliptical pillars (see Figure S2B for details on height and diameter measurements) that have an indentation or dimple close to the coverslip. The tapering stems from differences in the rate of chemical etching along the pillar while the dimple is a residual from the photolithography step of the fabrication process.

**Figure 3:**
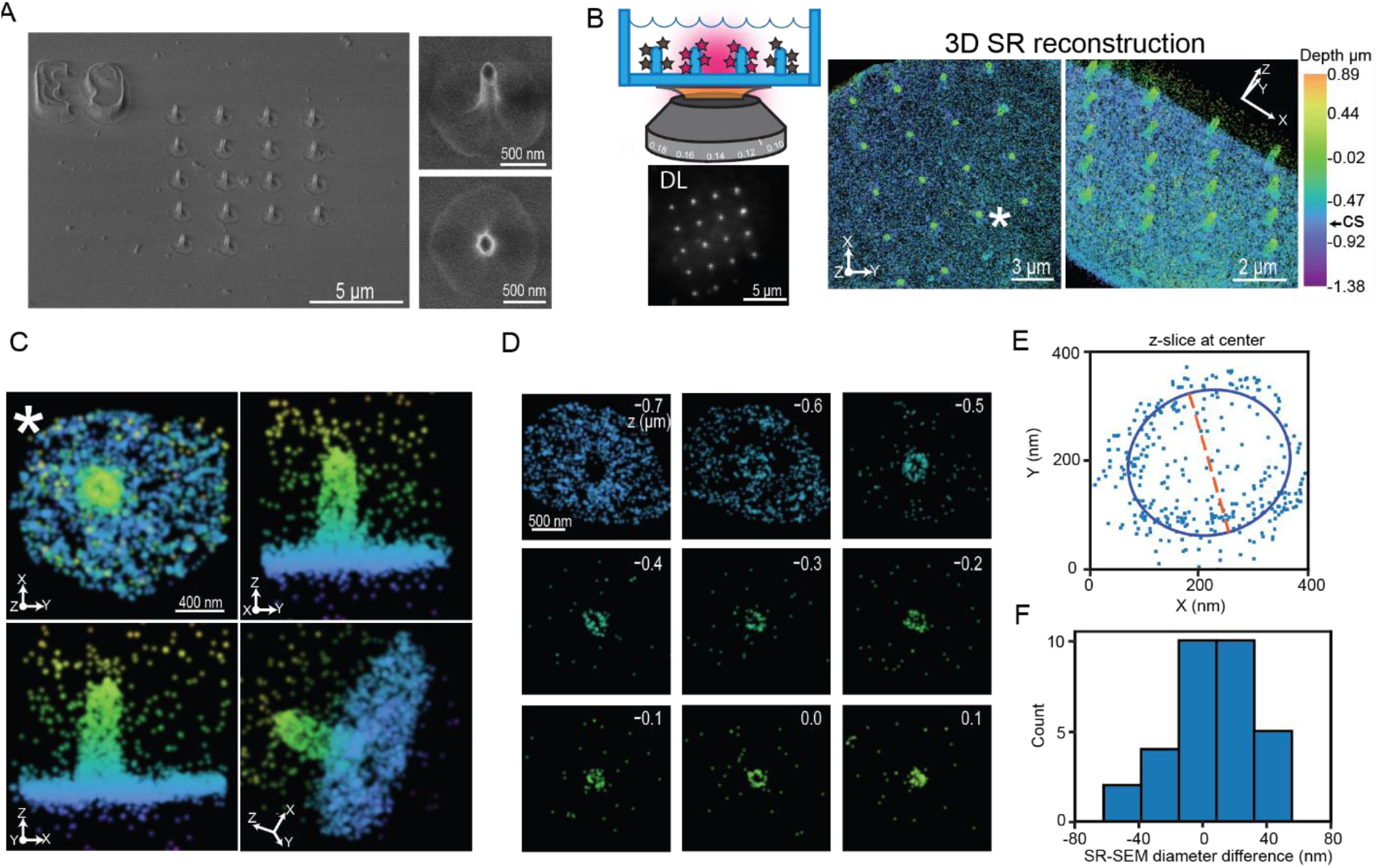
Comparison between 3D SR reconstructions of surface-labeled nanopillars and SEM images. (A) SEM images of a patterned array of nanopillars. The reference marker (E9) is clearly visible in the top left image of the nanopillars with a 30° tilt. The magnified images to the right of the image show an individual nanopillar; top with the tilt and bottom a top-down view. The dimple at the coverslip and elliptical shape of the pillars are clearly visible. (B) Top left is a cartoon depiction demonstrating imaging surface-labeled nanopillars. Below, the bright puncta in an array of surface-labeled nanopillars is visible in the DL image. Right: Two orientations of the 3D SR reconstructions show an array of nanopillars. Color encodes Z position. CS in calibration bar refers to the position of the coverslip. (C) Magnified images of an individual pillar (* in B) at various orientations. (D) 100 nm Z-slices of the pillar (C) from the bottom of the coverslip to the top of the nanopillar. Clear elliptical-shaped rings are visible. (E) 250 nm Z-slices of the 3D data shown as an XY projection at the center of an individual nanopillar fit to an ellipse. The orange dotted line is one axis extracted from the fit and may be compared to diameters extracted from SEM images. (F). Histogram depicts the difference in diameter between the 3D SR reconstructions and the SEM images of the nanopillars.

### Comparing SEM images and 3D SR reconstructions

With the fabricated and labeled nanopillars and the 3D DHPSF SR imaging in hand, the two imaging modalities may be directly compared as shown in Figure 3. Figure 3B depicts the DL and 3D SR reconstructions of the same nanopillar array in 3A. The DL image shows bright spots in the same array pattern as the SEM image. As the molecules along the shaft of the fluorescently labeled 3D nanopillar pillar will all project onto a 2D image corresponding to the focal depth of ~700-800 nm, the apparent density of fluorophore at the nanopillars will be higher compared to the coverslip. Thus, the bright spots in the DL image correspond to the positions of labeled nanopillars.

At the right of Figure 3B, the surface-labeled nanopillars were then imaged with the DHPSF 3D microscope. Any emitters that were poorly localized (XY precisions > 30 nm and Z precisions >40 nm) were filtered and removed. Further, we merged molecules to correct for overcounting (Methods and Figure S3A). After filtering, the median XY localization precision is 12 nm while the median Z precision is 27 nm, both precisions an order of magnitude better than that possible in DL imaging (Figure S3B for histograms of localization precisions). The XY projection of the 3D SR reconstructions (Figure 3B first of two right images) show an array pattern of regions containing a high density of localizations showing the location of the nanopillars. The color scale encodes the Z position in the reconstruction. As the coverslip is also labeled, the teal color in the reconstruction indicates the location of the coverslip (see Figure S3C for an additional 3D SR reconstruction).

Critically, the array patterns of the nanopillars from the DL image and the 3D SR reconstructions are in good agreement with the same pattern from the SEM images. In addition, rotating the 3D SR reconstruction to show a 3D perspective (far right Figure 3B) shows cylindrical, pillar-like structures similar to that in the SEM images. Figure 3C shows a magnification of one specific pillar at four different orientations (pillar zoomed in marked with * in Figure 3B). Qualitatively, the 3D SR results are quite similar to the SEM image. The pillar is straight and does not have any distorting shapes. As only the outer surface of the nanopillars are labeled with fluorophores, XY projections at various Z positions should appear as hollow elliptical rings given sufficiently high 3D precision. Figure 3D depicts 100 nm Z-slices from the bottom to the top of the nanopillar illustrated in Figure 3C. For all these slices except at the top of the nanopillar, elliptical hollow rings are evident. As the top cap of the pillar is also labeled, the top slice, as expected, is not a hollow ring but an ellipse filled with localizations. These Z-slices further underscore the high image quality in our 3D SR reconstructions. Furthermore, the Z-slices at the bottom (Z = −0.7 microns) show a dense number of localizations that surround the pillars. At Z = −0.6 microns, we observe a dark void roughly ~1 um in diameter surrounding the labeled nanopillar. Interestingly, this dark void is surrounded by localizations that do not stem from the nanopillar. These localizations are from the coverslip, and the appearance of the dark void 100 nm away from the bottom arises due to the dimple found at the bottom of the nanopillar. The many qualitative observations described here establish that the nanopillars from the 3D SR reconstructions have the same features and shapes as the pillars in the SEM images.

To quantitatively verify the high image quality, we compared the measured diameters of the pillars in the 3D SR reconstructions and the SEM images. First, we extracted the diameter at the axial half-way point in the SEM images by inspection of the SEM micrograph. To extract the diameters in the 3D SR reconstructions, we found the center of the pillars along z, and then took a 250 nm thick Z slice at the center (see Methods and Figure S4A for details).We then fit the XY projection of the localizations in this slice to an ellipse (Figure 3E), and used the major and minor axes of the ellipse fit as estimations for the diameter. Only the diameter from the fit that was at the same orientation of the SEM image in Figure 3A was used. Using the SEM reference marker as a guide, we then took the difference between the diameters from the 3D SR reconstruction and the SEM image of the same pillar for all the pillars analyzed (Figure 3F, n=31 pillars analyzed). The mean difference was 8.0±4.7 nm (mean ± standard error of the mean), indicating the diameters of our 3D SR reconstruction do not differ strongly from the SEM measurement, now benchmarked by correlative imaging.

As the pillars taper, we hypothesized the density of 3D single-molecule localizations would decrease as a function of distance away from the coverslip. To test this, we extracted 50 nm thick Z-slices from the bottom of the coverslip to the top of the pillar (Figure 4A and Methods for more details). The 50 nm Z-slices over many pillars (n=16) were binned into a histogram (Figure 4B), which shows that as the distance away from the coverslip increases, the density of localizations decreases, as we expect. To verify the behavior of this distribution, we simulated nanopillars with single-molecule localizations randomly decorating the surface of the pillars (Fig. 4C). The extent of tapering, the diameters, and the heights of the nanopillars were similar to the experimentally measured dimensions (see Methods for details). Critically, the probability that a localization was found at a specific region of the pillar was determined by the surface area at that section of the pillar to mimic the expected local behavior of the surface attachment. The positions of the simulated localizations were distorted in the XY and Z direction by Gaussian kicks with sigmas equivalent to the median XY and Z localization precisions to simulate realistic experimental conditions.

**Figure 4:**
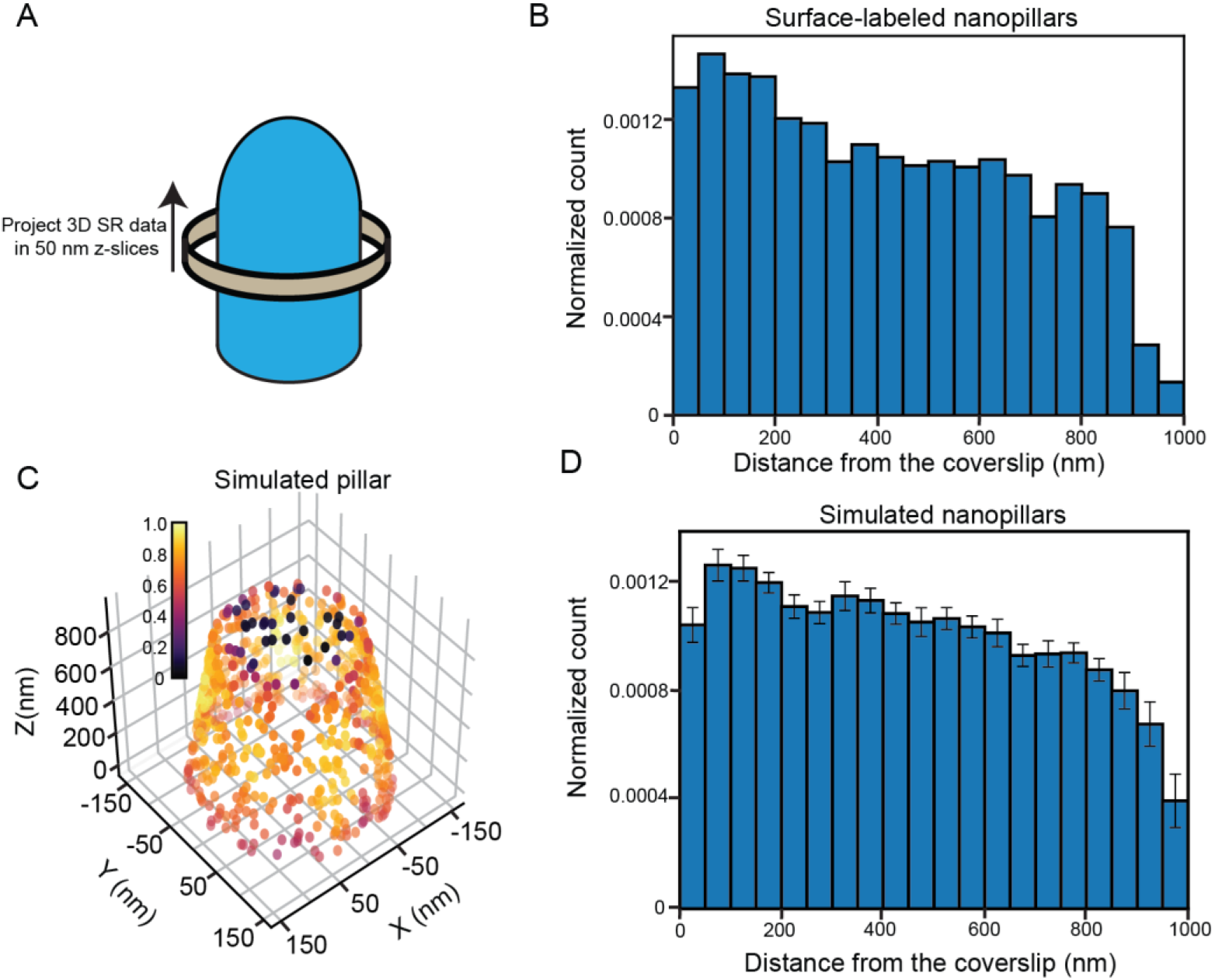
Quantification of density of molecules along nanopillars. (A). Cartoon depiction demonstration the protocol to extract the density of molecules along nanopillars form the 3D SR reconstructions of surface-labeled nanopillars. Localizations are projected onto the Z axis in 50 nm Z-slices from the bottom to the top of the nanopillar. (B). Compiled histogram of n=16 nanopillars demonstrating how the density varies along the nanopillar. As the pillars taper, we see the density decrease closer to the top. (C). 3D density plot of a simulated tapering nanopillar. Each localization is colored based on the local density. (D). Compiled histogram of n=16 simulated nanopillars. Distribution very similar to the experimental distribution in B. Values in bins of histogram are mean ± standard error of the mean (SEM).

Figure 4C depicts a 3D scatter plot of simulated nanopillar molecule localizations (see Figure S4B for an additional orientation) where the color encodes the number of nearest neighbors to each point. Qualitatively, the 3D scatter plot reveals that the number of localizations decreases closer to the top for the simulated nanopillar, similar to the experimental nanopillars. We then extracted 50 nm Z-slices along the pillar using the same methodology as the experimental measurement. Compiling the Z-slices over 16 simulated pillars (same number of experimental pillars used in this analysis) and binning the data into a histogram leads to the distribution in Figure 4D (see Figure S4C for distribution of simulated pillars without Gaussian kicks added). The shape of the distribution of the simulated pillars very closely resembles the shape of the distribution from the experimental data. This analysis verifies that the pillars taper with the measured localization density decreasing in an expected manner.

### Cell membrane-nanopillar interactions revealed from 3D SR reconstructions

Turning to cellular imaging, we first applied our method on a simple cellular system: the plasma membrane itself. Previous FIB-SEM and TEM approaches have imaged the membrane-nanopillar interface to reveal that the membrane can wrap around the pillars tightly^12, 13^. Thus, unlike proteins which may exhibit biologically more complex behavior near the nanopillars, the membrane is a relatively simple cellular system to first demonstrate that our 3D SR method is applicable to imaging the nano-bio interface in cells.

To label the membrane, we overexpressed a transmembrane domain of platelet-derived growth factor receptor (PDGFR) linked to a SNAP tag (Methods) in U-2 OS cells that were seeded onto the nanopillar substrate. The SNAP tag reaction was then used to attach AF647 with low nonspecific binding (see Figure S5A for controls). With this approach, the transmembrane domain is a single alpha helix that simply serves to anchor the fluorophore to the membrane. DL images (Figure 5A) show a labeled membrane with bright spots that are the nanopillars in the expected array pattern. After 3D DHPSF imaging, post-processing, filtering, and merging the data, the XY projection of the 3D SR reconstruction (Figure 5A, middle and right columns) also shows highly dense regions of localizations in an array corresponding to the locations of the nanopillars (median XY precision: 10 nm and median Z precision: 20 nm, see Figure S5B for distributions). This allows us to conclude that the area surrounding the nanopillar is the cellular membrane. As we image near the growing edge of the adhering cell whose membrane spreads on the substrate, we can observe many membrane protrusions. Projecting the reconstruction at a different orientation, the nanopillars appear as cylindrical pillar like structures, as expected (Figure 5A, see Figure S5C for an additional 3D SR reconstruction).

**Figure 5:**
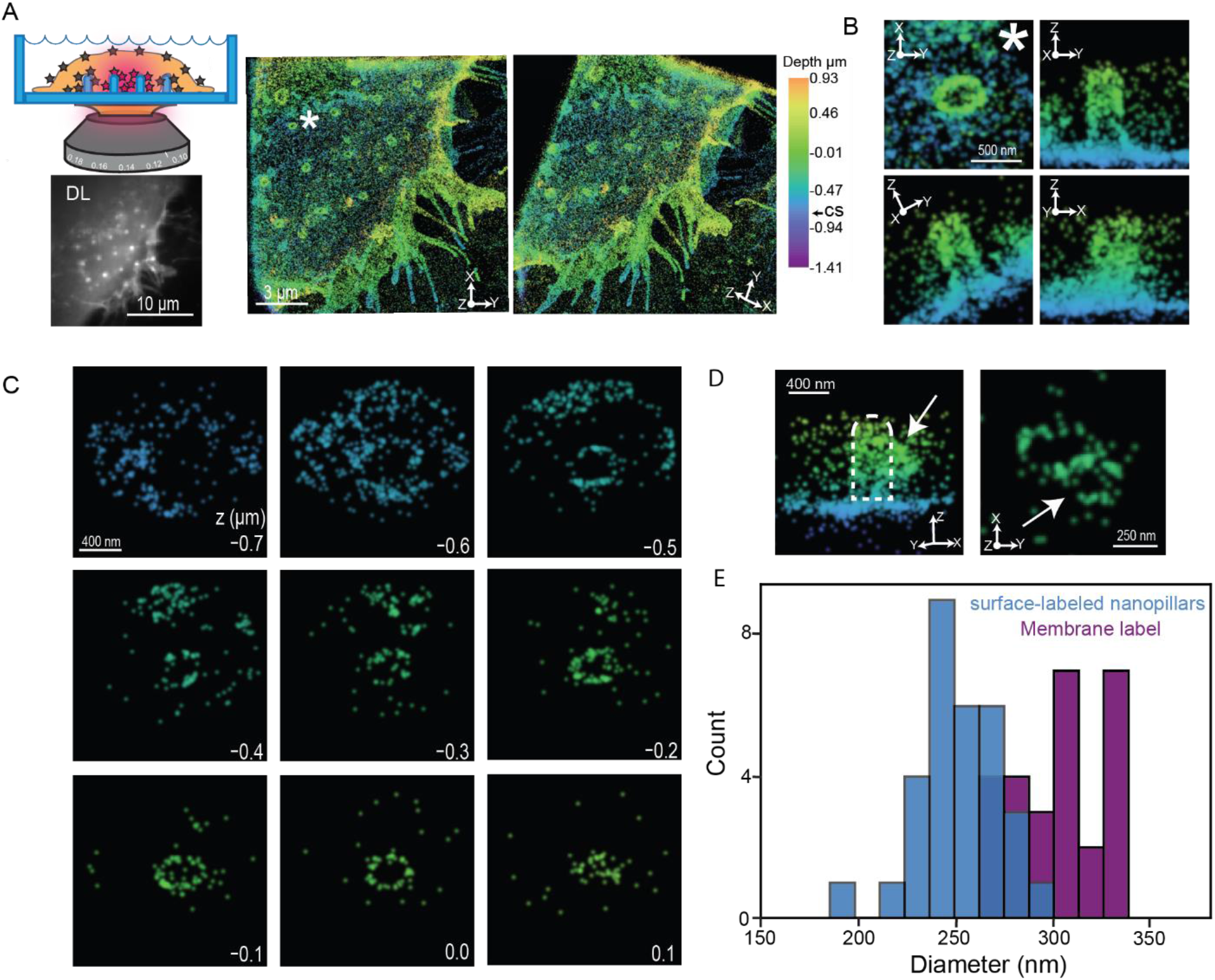
3D SR reconstructions of transmembrane-labeled cells. (A) Top left depicts a cartoon where we image the membrane at the nanopillars. Below the cartoon, the DL image shows bright puncta corresponding to the positions of the nanopillars in the DL image. The two 3D SR reconstructions at the right at different orientations reveal the membrane wrapping around the nanopillar array. Color encodes Z position and CS refers to the coverslip axial position. (B) Magnified images of a single pillar (* in A) at various orientations. The membrane is visibly wrapping around this pillar. (C) 100 nm Z-slices from the bottom to the top of the nanopillar in C. Bottom right values indicate Z position. Clear hollow elliptical rings are visible in the slices. (D) Two orientations of an individual pillar with a possible endocytosis event at the pillar. The white arrow points to the possible vesicle in both orientations. (E) Diameters at the center of the nanopillars extracted from surface-labeled and transmembrane labeled 3D SR reconstructions. The distribution from the transmembrane labeled cells is shifted towards higher values.

Figure 5B shows various perspective orientations of an individual pillar in the 3D reconstruction from 5A (denoted by the * in Figure 5A). We clearly see that the membrane wraps around the pillar tightly as it appears contiguous with the nanopillar. The membrane imaging also shows that the nanopillar is elliptical in cross section and is perpendicular to the surface. To benchmark the image quality of the reconstructions, we extracted 100 nm Z-slices from the bottom to the top of the membrane-wrapped nanopillar (Figure 5C). Similar to the surface-labeled nanopillar reconstructions, with sufficiently high 3D precision, the Z-slices appear as hollow elliptical rings as we labeled the membrane surrounding the outer surface of the pillar. Figure 5C also shows that the hollow elliptical rings change at the top cap region where we observe an ellipse filled with localizations. These Z-slices confirm the capability to observe membrane-labeled cells hugging the nanopillars.

Interestingly, for several nanopillars, we observed a bulge-like feature at certain Z positions along the pillar. Figure 3D (white arrow on left panel) shows one such example where a bulge is prominently protruding out from the membrane wrapped nanopillar. A 100 nm Z slice of the image on the right shows a hollow elliptical ring, the membrane wrapped-nanopillar, attached to another hollow ring. We hypothesize this attached hollow ring may be a vesicle forming and budding off from the nanopillar. Vesicles have previously been observed to bud off nanopillars from TEM images^12^, but observing such an effect is not easily possible with 2D DL imaging approaches.

Next, we probed how tightly the membrane wraps around the nanopillars. In previous work, some investigators have assumed that the membrane wraps tightly around the nanopillar, so that measurement of the membrane diameter is equivalent to the diameter of the nanopillar itself^11, 12^. Those studies have probed the influence of membrane curvature on the cell and curvature-sensing proteins indirectly by using nanopillar diameter as a proxy. In fact, studying how the membrane wraps around the nanopillars with EM imaging can be extremely challenging. For instance, while FIB-SEM methods have investigated the interaction between the membrane and nanopillars^13^, successful FIB-SEM imaging involves cross sectioning the interface between a hard (pillars) and soft (cell) materials. Cross sectioning between materials of different physical properties may induce artifacts. With the 3D SR method presented here, a fluorescence imaging technique can now be applied to directly probe the difference in the measured diameters between the nanopillar and the membrane.

To compare and extract the diameters by the two approaches, similar to Figures 3E and 3F, we found the center axial positions of the nanopillar regions in our surface-labeled and membrane-labeled 3D SR reconstructions. From this location, a 250 nm Z-slice was projected onto the XY plane and fit to an ellipse. In our previous analysis, we extracted a diameter derived from a single axis of the fit to match the SEM orientation (see Methods for more details). Here, we now simply average the diameters extracted from the two axes of the fit. The averaged diameters over many nanopillars (n=31 for surface-labeled nanopillars and n=27 for membrane label) are shown in the histogram of Figure 5E. The distributions show that the membrane labeled distribution is shifted towards larger diameters. The membrane labeled reconstructions have diameters of 302 ± 4.2 nm (mean±SEM) while the surface-labeled nanopillars have diameters of 250.7 ± 3.9 nm. This roughly 50 nm difference suggests ~25 nm gap distance between the cell membrane and the nanopillar surface, which agrees very well with previous measurements from FIB-SEM images^13^. Thus, our 3D SR method allows for relatively fast and simple imaging of the membrane at the nano-bio interface and illustrates that the difference between the diameters of the membrane and that of the underlying nanopillar can be observed.

### 3D SR reconstructions find that AP-2 distributes away from the nanopillar base

With successful application of our 3D SR method to the labeled plasma membrane, we next imaged a more complex biological system at the nano-bio interface: the spatial distribution of an intracellular protein AP-2. AP-2 is a multi-subunit protein complex that is s critical adaptor protein for clathrin-mediated endocytosis^46–48^. AP-2 has previously been shown to accumulate near nanopillars fabricated with high degrees of curvature^12^. To image AP-2, we used primary and secondary antibodies (see Methods and Figure S6A for controls) where the secondary has a blinking AF647 dye attached.

Figure 6A shows the DL image of AP-2 labeled cells grown on the quartz nanopillars. Similar to our previous results, the bright puncta in a specific patterned array show the location of the nanopillar and reveals that AP-2 accumulates at the nanopillars. The bright puncta are resolved as elliptical rings as shown in the XY projection of the 3D SR reconstruction in Figure 6A, middle image, and in the magnified images (see Figure S6B for additional reconstructions). In addition, we observe unstructured features colored with teal in the reconstruction. The magnified image (Figure 6A) shows an example of one unstructured feature more clearly. These structures, also termed plaques, form on the cellular surface next to the coverslip and have been observed in previous SR experiments imaging AP-2^49^. Thus, these plaques give us a reference to the coverslip position.

**Figure 6:**
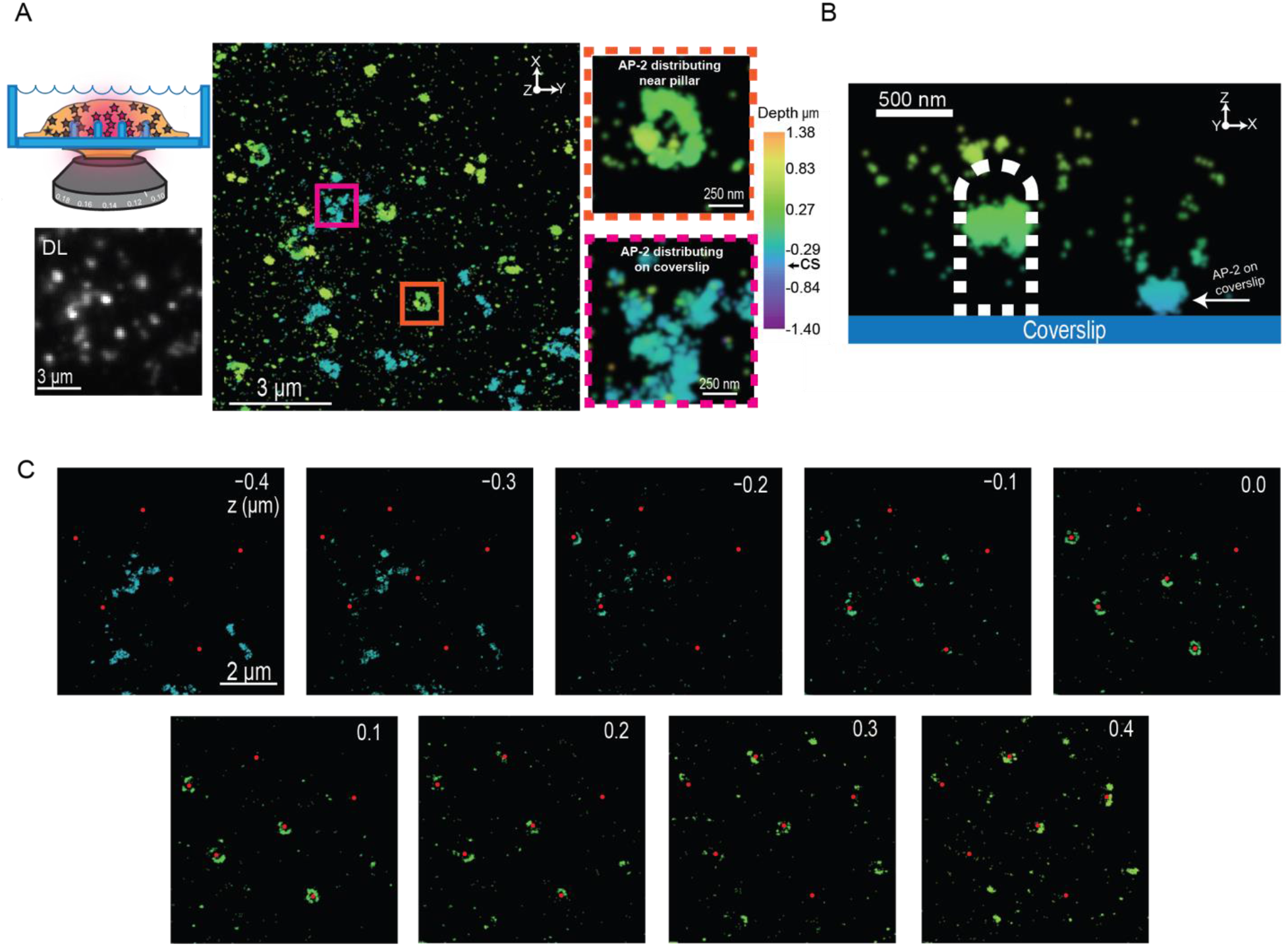
3D SR reconstructions of cells grown on nanopillars containing labeled AP-2 proteins. (A) Cartoon in the top left depicts imaging the cytoplasmic curvature-sensing protein AP-2 at the nano-bio interface. Below the cartoon, we see bright puncta corresponding to the positions of the nanopillars in the DL image. The 3D SR reconstruction to the right shows a large field, and AP-2 distributing around the pillars (orange inset) and also on the coverslip (magenta inset). Z position encoded by color and CS refers to position of coverslip in calibration bar. (B). XZ projection of an individual pillar. Dotted white line was manually drawn to indicate pillar shape and location. White arrow points to AP-2 distributing on the coverslip.AP-2 on the pillar appears to distribute away from the pillar bottom. (C) 100 nm Z-slices from the bottom to the top of the nanopillars. Red dots indicate positions of the nanopillar. AP-2 protein appear to form sectors of rings near the pillars and with few localizations at the pillar bottom (coverslip).

Strikingly, AP2 does not distribute uniformly on the surface of nanopillars like the membrane marker. The rings at the location of the nanopillars do not have AP-2 localizations encoded in the color teal, indicating that AP-2 does not localize around or near the bottom of the pillar. Figure 6B is a XZ projection of a region close to one nanopillar demonstrating the behavior of AP-2 along the pillar. Adjacent to the pillar, there is plaque locating the coverslip. However, on the pillar (depicted with the dotted white lines), we observe that the majority of AP-2 are distributed at a higher axial position on the nanopillar compared to the position of the plaque.

Figure 6C further confirms this result by showing 100 nm Z-slices from the bottom to the top of various nanopillars, where the red dots indicate the positions of the nanopillars. Close to the bottom (z=−0.4 to −0.2 μm), we observe very few AP-2 molecules distributed at the pillars. Instead, we observe the plaques that adhere to the coverslip. As the distance away from the coverslip increases, AP-2 molecules start to localize on the nanopillars forming rings and sectors of rings. Again, 3D SR imaging at the nano-bio interface shows features not observable with conventional 2D DL imaging. Only with sufficient resolution and 3D information can we observe that AP-2 prefers to distribute away from the coverslip at higher Z-positions along the nanopillar shaft. This result will be explored in more quantitative detail below.

### 3D SR reconstructions of actin molecules, fibers, and bundles distributing at the nanopillars

Next, we investigated the nanoscale distribution of actin molecules at the nano-bio interface. Actin is a well-known cytoskeletal protein with many functions, for example polymerization to form fibers that are critical for processes such as cell motility^50^, cell division^51^, and clathrin-mediated endocytosis^52^. In addition, the polymerization of actin fiber is curvature sensitive and has been shown to reorganize upon changes in membrane curvature^11^. To label actin, we used phalloidin, a small molecule that binds specifically to actin fibers, linked with AF647 (see Methods and Figure S7A for controls).

Figure 7A, left, shows the DL image of an actin-labeled cell. We observe bright puncta organized in an array corresponding to actin fibers distributing at the nanopillars as previously reported. We also observe actin fibers throughout the cell. Figure 7A (middle and right) shows an XY projection of the 3D SR reconstruction (see Figure S7B for an additional reconstruction). We clearly observe actin fibers that are located at various Z positions in the reconstruction. In addition, we see regions where actin is accumulating around the nanopillars. To observe these features more clearly, Figure 7A, right, also shows a 300 nm thick Z slice at the coverslip. The white arrows depict the actin fibers and the regions where actin accumulates around the nanopillars.

**Figure 7:**
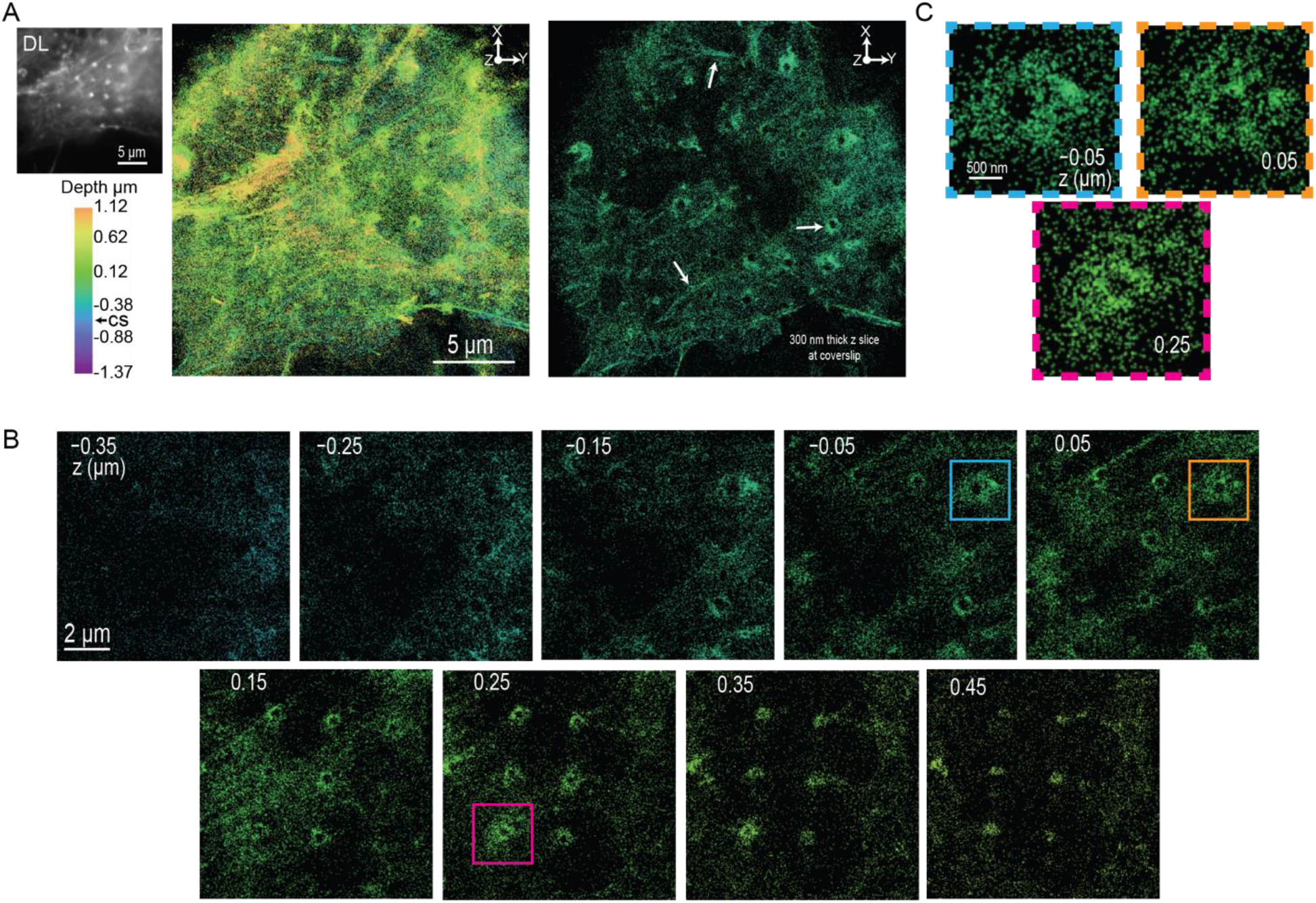
3D SR reconstructions of labeled actin in cells incubated on nanopillars. (A) DL image (top left) shows a cell where the bright puncta correspond to regions of the nanopillars. The XY projection of the 3D SR reconstruction (middle) shows nanopillar regions and actin fibers. The 300 nm Z-slice close to the coverslip (right) more clearly shows the nanopillar regions and actin fibers (white arrows). (B) 100 nm Z-slices from the bottom to the top of the nanopillars. We observe the actin fibers and elliptical hollow rings at the nanopillars. (C) Magnified images of actin at specific pillars and Z-positions shown in B (magenta, teal, and orange boxes). The actin at these locations appears to form hair-like structures that are difficult to observe with DL imaging.

We extracted 100 nm Z-slices from the 3D SR data from the bottom to the top of various nanopillars to more clearly visualize the behavior of actin around the pillars. As expected, in these Z-slices, we see hollow rings that are generally surrounded by actin molecules. Nearby actin fibers are prominently seen located by the pillars at Z = 0.15 microns. Strikingly, a few of the nanopillars appear to be surrounded by a hair-like structure as well. Potentially this hair-like structure arises from actin bundles accumulating at the nanopillars or from fibers that wrap around the pillars (Figure 7C). These hair-like features on the pillars are large (1-2 microns) but the fine features and are much more challenging to clearly observe in the DL image because they seem to occur only at certain Z-positions.

### AP-2 molecules distribute along nanopillars at increased 1/R membrane curvature regions

The 3D SR reconstructions of the membrane, AP-2, and actin, all reveal features that now may be quantified. We consider the density of molecules along the pillars and compare the behavior of the various labels and biomolecules. For instance, AP-2 appears to distribute closer to the top of the pillars. In contrast, the membrane appears to have a more homogenous density of molecules along the entire pillar. To quantify the density of localizations for membrane, AP-2, and actin reconstructions, we projected 3D localizations of the molecules surrounding the nanopillars into 50 nm thick Z-slices, similar to our previous analysis with the surface-labeled nanopillars (Figure 4A depicts a cartoon describing the analysis). For each of the labeled targets, the 50 nm Z-slices over many pillars were binned into histograms.

Figure 8A shows the distribution of molecules along the nanopillars for the membrane (n=37 pillars analyzed). Given that the pillars taper, the surface area available to the membrane will thereby decrease and thus, we would expect that the number of molecules would gradually decrease towards the top of the pillar, maximizing near the bottom of the pillar. While the density of molecules does gradually decrease closer to the top of the nanopillar, we observe that the number of molecules near the very bottom (Z = 0-100 nm) is low and progressively increases to a maximum around Z = 200 nm. This result is consistent with previous findings^53^ where the membrane did not fully wrap near the bottom of the nanopillar. Instead, the membrane rises from flat regions near the coverslip until the membrane encounters a nanopillar where it then wraps around the pillar.

**Figure 8:**
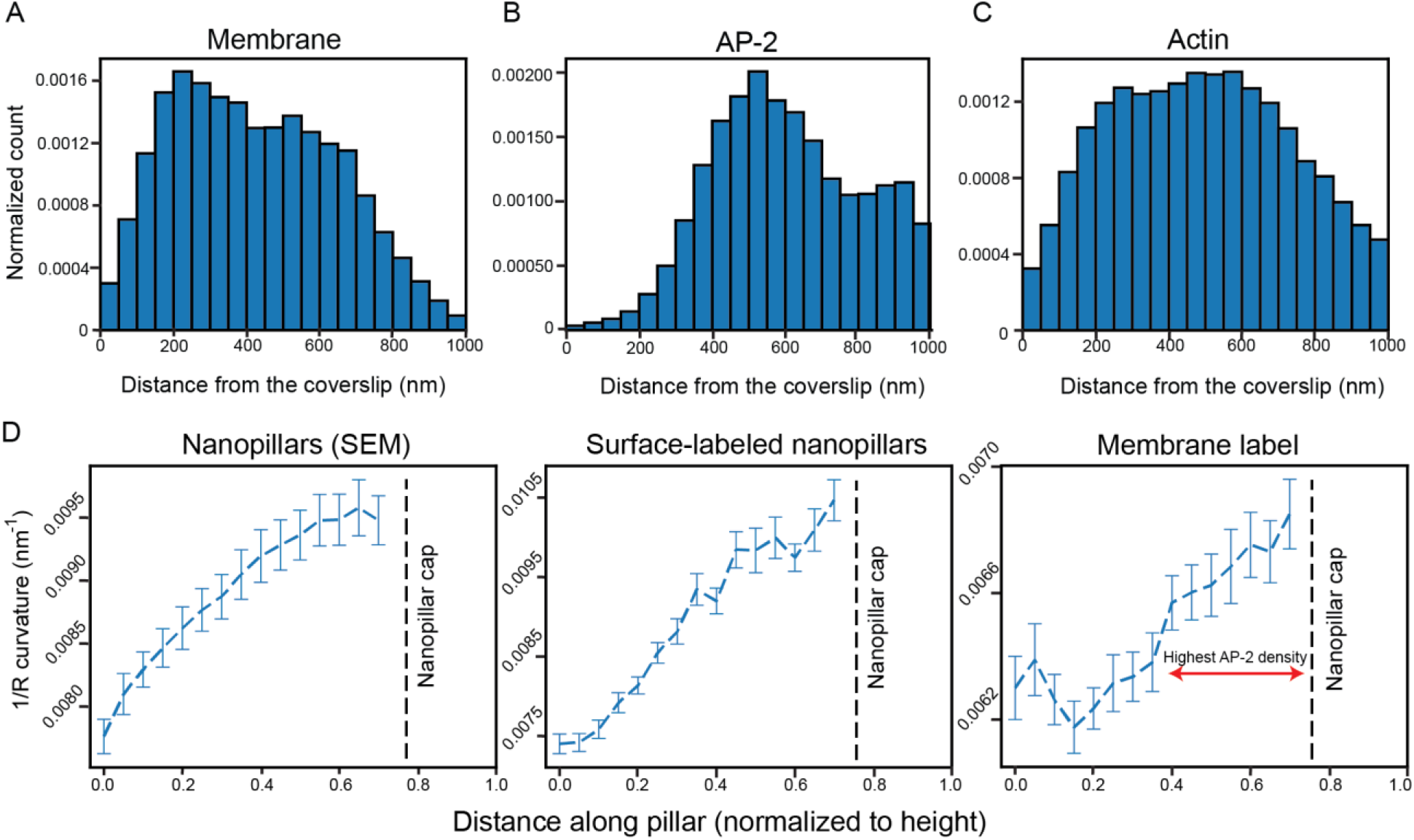
Quantifying molecular positions and the variation of 1/R curvature along the nanopillars. (A) Distribution of molecules along nanopillars for transmembrane labeled cells quantified by projecting localizations in 50 nm Z-slices along pillar. We observe very few molecules near the bottom of the pillar and the distribution peaks around Z = 200 nm. As the pillars taper, the density decreases closer to the top. (B). Distribution of molecules along nanopillars for AP-2. The density peaks around 500 nm and begins to decrease. Very few molecules are observed from Z = 0 to Z = 200 nm. (C) Distribution of molecules along nanopillars for actin. Similar to the membrane distribution, very few molecules are found near the coverslip. The distributions seem to peak near the middle and the density decreases closer to the top. (D) The 1/R curvature along the nanopillars is calculated using the SEM images (left), the surface-labeled 3D SR reconstructions (middle), and the transmembrane labeled 3D SR reconstructions (right). The curves for the SEM and surface-labeled nanopillar panels are similar. The double red arrows on the membrane curvature panel shows the regions where AP-2 preferentially distributes at the nanopillar. The dotted black line shows where the nanopillar cap is located and 1/R curvature was not analyzed in this region. Values in curves are mean ± SEM.

Figure 8B is the histogram for the AP-2 labeled cells (n=49 pillars analyzed). The distribution significantly differs from the membrane-labeled distribution and reflects the behavior we observed in the 3D SR reconstructions of AP-2 labeled cells (Figure 6). We observe that the density is very low from Z = 0 to Z = 200 nm. From Z = 200 nm, the density gradually rises until it reaches a maximum at Z = 500 nm. Further up the nanopillar, we would expect that the density of molecules might decrease above Z = 700 nm as the cap is approached. While the density does indeed decrease, strikingly, the density decreases at a rate much smaller than the rate found in the distribution derived from membrane labeled cells. This result reveals that AP-2 does not homogenously distribute on the membrane along nanopillars. Instead, AP-2 appears to prefer the middle and top of rather than the bottom of the nanopillars.

Figure 8C depicts the histogram for actin labeled cells (n=27 pillars analyzed). Similar to the membrane, the density of molecules from Z = 0 to Z = 100 nm is low. The density gradually increases until a maximum is reached around Z = 450 nm. The density of molecules decreases from the maximum, although, similar to the AP-2 labeled cells, the rate of decrease is much slower compared to the rate for membrane-labeled cells in Figure 8A. In general, the actin and membrane distributions are similar except the rate decreases closer to the top of the nanopillars.

As the nanopillars taper, their 2-dimensional curvature, the reciprocal of the radius of a cross section at fixed Z (which we term 1/R curvature), increases from the bottom to the top. (For the nearly cylindrical structure of the pillar shaft, the axial curvature is approximately zero.) Thus, we hypothesized that it is this strong variation in 1/R curvature along the pillars that drives AP-2 to preferentially accumulate at regions away from the bottom of the nanopillar. To first assess the extent of tapering, we extracted the diameter along the pillars from the SEM images to calculate the 1/R curvature (see Methods and Figure S8A for additional details). For simplicity, we only extracted the diameters at the orientation shown in the side view magnification in Figure 3A. As the very top or the cap of the nanopillar is approximately shaped as a hemisphere, the 2D 1/R curvature metric is not well suited for the 3D surface at the top. Thus, we have excluded calculations of 1/R curvature near the top of the nanopillar for all our results.

Figure 8D shows how the 1/R curvature changes along the pillar. We clearly observe that as the Z position increases, the curvature increases as well. However, as we described earlier, the pillar curvature is not exactly equivalent to the membrane curvature. Before extracting the membrane curvature, as a positive control, we first extracted the 1/R curvature along the pillar for our surface-labeled 3D SR reconstructions to compare to the 1/R curvature measurements from the SEM images (left and middle panels). To extract the curvature, we first measured the diameters of the pillars by fitting ellipses to projected 250 nm Z-slices that were centered at various axial points along the pillar (see Methods and Figure S8B) similar to our analysis above. From the ellipse fit, we extracted the diameter that was the same diameter measured in the side view SEM images in Figure 3A. Using the diameters and repeating this analysis procedure over many pillars, the 1/R curvature was calculated and plotted as a function of position along the pillar in Figure 8D (middle). We can clearly see that the plots and absolute values of 1/R curvature from the SEM images and SR reconstructions are reasonably similar underscoring the good correspondence between the two methods.

Next, we calculated the 1/R plasma membrane curvature at the nano-bio interface using the same analysis protocol used to calculate the plot in Figure 8D with slight modification. Instead of selecting for a diameter from either the major or minor axes of the ellipse fit, the diameters were simply averaged to calculate the 1/R curvature for the membrane. Figure 8D (right) reveals how the membrane curvature varies along the pillar, increasing near the top due to the tapering. In addition, the 1/R membrane curvature is also larger than the 1/R curvature of the nanopillars themselves, as expected from the separation between the nanopillar and the membrane in Figure 3E. Critically, the membrane curvature plot reveals that AP-2, in particular, does not appear to preferentially accumulate at low degrees of membrane curvature near the bottom of the pillar (Figure 8B). Only after the membrane curvature increases its value above a specific threshold (shown by the red line in the right panel), we observe AP-2 beginning to preferentially accumulate at the nanopillar.

### Positive Gaussian curvature at pillar caps increases relative density of AP-2 and actin molecules

The analysis presented above clearly reveals that increased 1/R curvature leads to a higher density of AP-2 molecules at higher axial positions along the pillar. While the curvature analysis at the cap of the nanopillar was excluded, the distributions in Figure 8B and 8C near the cap differ from the membrane distribution. Thus, we investigated how AP-2 and actin behave at the cap more closely. The behavior near the cap, however, is obfuscated by variations in the membrane surface area, which may be regarded as the property that is locally sensed by these proteins. For instance, closer to the top of the pillar, the membrane surface area is smaller and thus, the density of molecules will decrease. To account for these effects, we normalized the distributions in Figures 8B and 8C by the membrane surface area found at each Z position of the histograms. To do this, we first assumed that the number of molecules in each Z position in the membrane distribution in Figure 8A is proportional to the membrane surface which can also be calculated at the cap region. Then, each Z position along the pillar in the AP-2 and actin distributions was divided by the molecular count found at each Z position in the membrane distribution.

The distribution in Figure 9A depicts the normalized distribution for AP-2 along the entire pillar. Similar to the unnormalized distribution the relative density of AP-2 molecules is close to zero from z= 0 to z= 200 nm, rises to a local maximum at z= 500 nm, and then plateaus until Z = 750 nm with the relative density of molecules being nearly double the density found in the membrane labeled cells. Strikingly, after Z = 750 nm, the distribution rapidly increases and maximizes at the cap of the nanopillar where the relative density is now nearly 8 times larger than the density found in the membrane. In addition, Figure 9B depicts the normalized histogram for actin which is similar, but with variations. First, from Z = 0 nm to Z = 750 nm, the density of actin molecules is roughly equivalent to the density of membrane molecules at these positions. However, past 750 nm, the relative density increases sharply until it maximizes at the top of the pillar. At the cap of the pillar, density of actin molecules is five times greater than the density of membrane molecules.

**Figure 9:**
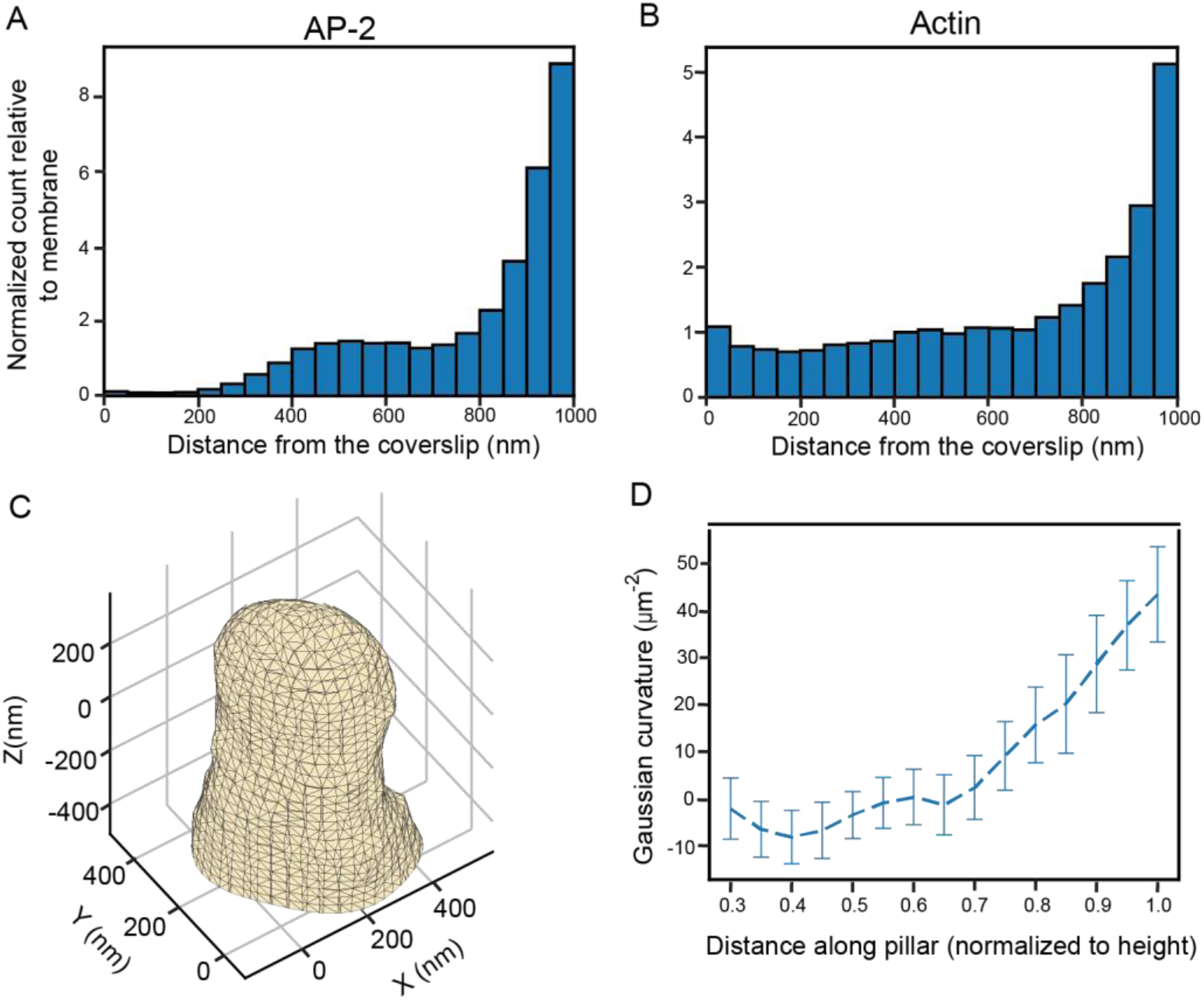
Quantifying density of molecules normalized to membrane surface area and Gaussian curvature analysis along nanopillars. (A) Distribution of molecules of AP-2 in cells along nanopillars normalized to membrane surface area. We observe very few relative molecules near the bottom of the coverslip. The density is nearly twice that density found at the membrane from Z = 400 to Z = 800 nm. Near the top of the pillars, the relative density increases substantially. (B). Distribution of actin molecules in cells near nanopillars normalized to membrane surface area. The density relative to the membrane is nearly constant until around Z =800 nm. The density increases greatly after this point. (C). Representative mesh surface derived from 3D localization data from the transmembrane labeled 3D SR reconstruction. (D). Gaussian curvature along the nanopillar. Curvature is nearly 0 until 0.8 where it increases to relatively large positive values. Values are mean ± SEM.

The rapid increase in the density of AP-2 and actin molecules at the cap of the nanopillar indicates that the 3D shape of the pillar near the top influences the behavior of these proteins. We hypothesize that since the hemielliptical shape of the cap induces positive Gaussian curvature, it is this property that influences the behavior of AP-2 and actin near the cap region. Gaussian curvature is a mathematical metric that is used to describe the curvature of 3D surfaces (see Methods). Spherical objects such as spheres and ellipsoids have positive Gaussian curvature while saddle surfaces feature negative Gaussian curvature. The surfaces of cylinders have zero Gaussian curvature. For our nanopillars, we would expect that as the body of the pillar is a truncated cone, the Gaussian curvature is 0 along the shaft of the nanopillar while the hemielliptical cap has positive Gaussian curvature.

We used approaches previously described to extract the Gaussian curvature along the nanopillars^45^. Briefly, the screened Poisson surface reconstruction^54^ algorithm was applied to the 3D localizations of the nanopillars to create a 3D triangulated surface mesh using the software MeshLab^55^ (Methods and Figure S9A). We created surfaces meshes using both the surface-labeled (n=15) and membrane labeled (n=14) 3D SR reconstructions. Figure 9C shows a surface mesh derived from the membrane labeled reconstructions (see S9B for surface meshes from surface labeled reconstructions). From the surface, we can clearly see a pillar structure that has a hemielliptical cap, as expected. The surface mesh additionally reflects the positions of the 3D localizations and is hollow (Figure S9C).

Gaussian curvature was then calculated along the nanopillar as described previously^56^. The Gaussian curvature values along the mesh was projected into Z slices spaced equidistant along the pillar. The average Gaussian curvature in slices along the pillar was then calculated. As expected, the Gaussian curvature from the meshes of the surface labeled pillars (Figure S9D) is zero along the body and increases to positive values at the cap. In addition, we analyzed the Gaussian curvature from our simulated nanopillars and find (n=15, Figure S9E) similar behavior to the results from the surface-labeled nanopillars, further confirming our hypothesis that the Gaussian curvature would be highly positive near the cap. Finally, Figure 9D reveals the variation in Gaussian curvature from surfaces of the membrane labeled reconstructions. We clearly see close to zero Gaussian curvature along the body and highly positive Gaussian curvature near the cap. The highly positive values at the cap correlate well with the increased relative density of AP-2 and actin molecules near the cap in Figures 9A and 9B. This result indicates that the positive Gaussian curvature may be the driver which increases the density of curvature-sensing proteins.

Our results show that 2D and 3D variants of membrane curvature metrics, 1/R curvature and Gaussian curvature, may influence the behavior of specific curvature-sensing proteins. Previous studies have assumed that the 1/R curvature is critical for the behavior of curvature-sensing proteins at the nano-bio interface. Here, we show that the shape of the pillar is also critical for the behavior of these proteins. The positive Gaussian curvature at the top induces changes that may increase the density and behavior of the curvature-sensing proteins at the cap. Thus, future studies should systematically study the effects of 1/R and Gaussian curvature separately for various curvature-sensing proteins to understand the contribution of each variant of curvature. This conclusion is enabled by using our described 3D SR method that allows probing 3D localizations of specific proteins at the nanoscale on nanofabricated quartz surfaces.

## Conclusion

We have shown that having 3D SR information about the positions of the plasma membrane and various curvature-sensing proteins can lead to new insight about the cellular behaviors near nanopillars seeking to define a particular nano-bio interface. This arises from several factors: first the higher spatial resolution, second the 3D character of the information, and finally, since the acquisition method involves measuring the localization of many single molecules sampling the object of interest, it is then possible to apply powerful statistical methods from point-based image analysis to extract further insight. While previous studies were mostly focused on the importance of 1/R curvature effects, here, we show distinct effects of 1/R curvature vs. Gaussian curvature in affecting protein localizations. This ability can then be applied in future studies in combination with assessment of cellular signaling changes. By directly demonstrating the power of the technique on realistic cellular imaging problems, we expect this approach to be widely applicable to other cellular imaging problems where nanoscale objects nearby drive the cellular response and behavior.

## Methods

### Nanopillar fabrication

Quartz substrates were ordered from Technical Glass Products with dimension of 1 inch x 1 inch and 200 um thickness (Technical Glass Product 1×1×0.2). The substrates were cleaned by isopropyl alcohol and sonication to remove any surface particles. The substrates were then dehydrated at 180C on a hotplate for two minutes and then coated with hexamethyldisilazane (HMDS) to promote resist adhesion. Arrays of nanopillars were fabricated on these prepared substrates using a photolithography and wet etching technique. In the photolithography step, a positive photoresist (Shipley 3612) was spin coated on the substrate and exposed by a mask-less aligner (Heidelberg MLA-150) to form arrays of circular holes. A chromium mask was later deposited onto the patterned substrates using electron-beam evaporation, and excess resist was removed by acetone, resulting in arrays of circular chromium disks for pillar fabrication. Vertical pillars were subsequently fabricated by anisotropic reactive ion etching with a mixture of CHF3, C4F8, and Ar (Versaline LL-ICP Oxford etcher) to produce a tapering vertical profile. The substrates were then immersed in Chromium Etchant 1020 (Transene) to remove the chrome mask. Nanopillars were subsequently modified through a wet etching technique to shrink the pillar diameters. The final nanopillar dimensions were precisely controlled by submerging the substrates in a Buffered Oxide Etchant 20:1 (Transene). The final nanopillars were characterized with scanning electron microscopy.

### SEM imaging of nanopillars

The SEM images were taken using a FEI Nova (NanoSEM 450). Since quartz is a non-conductive substrate, the imaging was operated with very low voltage (2kV). The images were taken with an Everhart-Thornley Detector in the Field-free mode at lower magnification, and in the immersion mode with Through-lens detector for high resolution imaging at high magnification.

### DNA vectors

The vector for the eGFP-FBP17 protein fusion, pEGFP-C1-FBP17, was a gift from Pietro De Camilli (Addgene plasmid # 22229). To construct the vector for cell surface tethered SnapTag expression, the DNA fragment encoding SnapTag is cloned into pDisplay (Invitrogen, V66020) between restriction sites BglII and SalI.

### Cell culture and transfection

U-2 OS cells (ATCC^®^ HTB-96™) were maintained at 37°C, 5% of CO_2_ atmosphere in complete cell culture medium (Dulbecco’s Modified Eagle’s Medium, DMEM) (Sigma-Aldrich, D6429) supplied with 10 % (v/v) Fetal Bovine Serum (FBS) (Sigma-Aldrich, F4135), 100 units/mL of penicillin and 100 μg/mL of streptomycin (Gibco, 15140122). For microscopic imaging, U-2 OS cells were detached using TrypLE Express enzyme (Gibco, 12604013) and plated on gelatin (Sigma-Aldrich, G9391)-coated nanopillar substrates in complete cell culture medium. Before cell plating, quartz nanopillar substrates were first treated by air plasma (Harrick Plasma) for 15 minutes and then incubated in phosphate-buffered saline (PBS) with 0.1 mg/mL poly-L-lysine (Sigma-Aldrich, P5899) for 1 hour. Afterward, the nanopillar substrates were washed with PBS three times and incubated in PBS with 0.5% (v/v) glutaraldehyde (Sigma-Aldrich, G6257). After further washing with PBS (3X), the substrates were then incubated in PBS with 0.5% gelatin (Sigma-Aldrich, G9391) for 1 hour at 37°C. The coated substrates were finally washed with PBS (3X) and treated with 1 mg/ml sodium borohydride (Sigma-Aldrich, 452882) in PBS for 5 minutes to eliminate autofluorescence.

For the expression of GFP-FBP17 and cell surface-tethered SnapTag, cells were transfected with DNA vectors by electroporation. U-2 OS cells were grown in 6-well plates (Corning, 353046). For transfection, one well of cells were detached from the culture plate using TrypLE Express enzyme and spun down at 300 relative centrifugal force (RCF) for 3 minutes. The supernatants were removed as completely as possible, leaving cell pellets which were then resuspended in electroporation mix containing 100 uL Electroporation buffer II (88mM KH2PO4 and 14 mM NaHCO3, pH 7.4), 2uL Electroporation buffer I (360mM ATP + 600mM MgCl2) and 1ug DNA vector. The electroporation was executed in a 2 mm-gap electroporation cuvette (Invitrogen, P45050) by Amaxa Nucleofector II following manufacturer’s protocol. The cells recovered from electroporation in 650 μL complete cell culture medium for 5 minutes at room temperature (RT) and were plated on nanopillar substrates. The cells were grown for 24 hours before next treatment.

### Labeling

#### Surface labeling of nanopillars

The nanopillar substrates were cleaned with 1M potassium hydroxide (Sigma-Aldrich, 221473) solution for 15 minutes at RT. The substrates were washed with nanopure water five times and air dried. The substrates were then treated with air plasma for 15 minutes and attached to plastic dishes, which have a hole punched in the bottom, with silicone sealant. The substrates were rinsed with anhydrous methanol (Sigma-Aldrich, 322415) and incubated in 2 ml of a mixture containing anhydrous methanol, glacial acetic acid (Sigma-Aldrich, 695092) and (3-Aminopropyl) triethoxysilane (Sigma-Aldrich, A3648) in a v:v:v ratio of 100:5:3, for 30 min. The substrates were washed with anhydrous methanol five times and then with nanopure water for three times. Afterward, the substrates were rinsed with 0.1M sodium bicarbonate (Sigma-Aldrich, S5761) solution (pH 8) and incubated in 0.1M sodium bicarbonate solution (pH 8) with 2 uM Alexa Fluor™ 647 NHS Ester (Invitrogen, A37573) for 30 minutes. The substrates were washed with PBS five times before imaging.

#### Membrane labeling

The pDislay-SnapTag transfected cells were cultured in complete cell culture medium for 24 hours before labeling. The cells were incubated in labeling solution, 5% CO_2_ -balanced complete cell culture medium with 5 μM SNAP-Surface Alexa Fluor 647 (NEB, S9136S), for 15 minutes at 37°C. Before being added to the cells, the dye and medium were mixed thoroughly by pipetting up and down ten times. The cells were then quickly washed with CO_2_ -balanced complete cell culture medium five times and fixed with 4% paraformaldehyde (PFA) (Sigma-Aldrich, 158127) in PBS for 15 minutes at RT. The samples were washed with PBS three times before imaging.

#### GFP nanobody labeling

Anti-GFP nanobody (Chromotek, gt-250) was diluted in 0.2 M sodium bicarbonate solution at pH 8.2, to a final concentration of ~60 uM. Alexa Fluor 647 NHS Ester stock solution (1mg/ml in DMSO) was added into diluted anti-GFP nanobody to a final dye concentration of ~120 uM. The mixture of nanobody and dye was incubated for 1 hours at 25°C. Then, free dyes were removed from the solution using a Zeba spin desalting column (Thermo Scientific, 89882). As the extinction coefficients are known, the concentrations of purified nanobody and conjugated AF647 dye were determined using light absorption at 280-nm wavelength and 647-nm wavelength, respectively. The degree of labeling (the average number of dye molecules per protein) was calculated to be 1.08.

#### Immunofluorescence labeling

The cells were fixed with 4% PFA in PBS for 15 minutes at RT. The fixed cells were washed with PBS three times and permeabilized with 0.1% triton-X (Sigma-Aldrich, T9284) in PBS for 15 minutes at RT. The samples were washed with PBS three times and then blocked in PBS with 5% (w/v) bovine serum albumin (BSA, Sigma-Aldrich A3059) overnight at 4°C.

To label GFP-FBP17, the samples were incubated in PBS with 5% BSA and 1 nM AF647-conjugated GFP nanobody for 2 hours at RT. The samples were then washed with PBS containing 0.1% triton-X and 5% BSA three times for 15 minutes each and PBS five times for 2 minutes each before imaging.

To label endogenous AP2 complexes, the samples were incubated in PBS with 5% BSA and Anti-alpha Adaptin primary antibody (Abcam, ab2730, 1:250) for 2 hours at RT, and washed with PBS containing 0.1% triton-X and 5% BSA three times for 15 minutes each. The samples were then incubated with goat anti-mouse secondary antibody, Alexa Fluor 647 (Invitrogen A32728, 1:500) in PBS with 5% BSA for 2 hours at RT. The samples were finally washed with 0.1% triton-X and 5% BSA in PBS five times for 5 minutes each, and with PBS three times before imaging.

#### Actin labeling with phalloidin

Cells seeded on chips were fixed with 4% PFA for 15 minutes and subsequently washed three times with PBS. Cells were incubated with 5% BSA in PBS for 1 hour. Following incubation, 330 nM of phalloidin conjugated AF647 (Cell Signaling Technology, 8940) was added to the solution for 15 minutes. Then, samples were washed once with PBS before imaging.

### Super resolution microscopy

#### Optical setup

All data and images were acquired using our custom built widefield double-helix PSF inverted microscope. We imaged our samples using a 1 W 647 nm continuous wave (CW) laser (MPB Communications) for 3D SR imaging and a 100 mW 641 nm CW (Coherent Cube) for 2D SR imaging and calibration of the silicone oil objective. In our setup, the 647 nm excitation laser was first passed through a clean-up excitation bandpass (631/36) filter (Semrock, FF01 -631/36-25) and then a quarter wave plate (Thorlabs,WPQSM05-633) for circular polarization. The size of the laser beam was then magnified twice using a two pairs of lenses before entering the backport of the microscope (Olympus IX71). A Köhler lens placed before the backport was used to focus the light at the back focal plane of the objective for widefield imaging. Inside the microscope, a dichroic mirror (Semrock, Di01-R405/488/561/635-25×36) was used to relay the light through the objective. The standard oil immersion objective (UPlanSAPo 100x/1.4 oil, Olympus) with immersion oil (Zeiss, 444960-0000-000) was used for imaging samples on glass substrates. The silicone oil immersion objective (UPlanSAPo 100x/1.35 silicone oil, Olympus) with silicone immersion oil (Olympus, Z-81114) was used for imaging samples on quartz substrates and for imaging nanopillar samples. The samples were mounted on a motorized XY stage (Physik Instrumente, U-780.DOS) and a precision XYZ piezo stage (Physik Instrumente, P-545.3C8). The emitted fluorescence light was collected using the objective, transmitted through the dichroic and then focused using the tube lens (f=180 mm) inside the microscope to the standard intermediate image plane position. The emitted fluorescence was relayed using two lenses (f=90 mm for both lenses) in a 4f optical configuration to access the Fourier plane of the microscope, enabling insertion of the double-helix phase mask for 3D SR imaging. The double-helix phase mask (emission wavelength 665 nm, diameter 2.8 mm, fabricated with fused silica, Double-Helix Optics LLC) was removed for DL imaging or 2D SR experiments. Emission filters (ET700/75 bandpass filter, Chroma, ZET647 notch filter, Chroma, 680/60 bandpass filter, Omega) for imaging with the 647 nm laser were placed in the 4f emission pathway to remove reflected laser light and Raman scattering. The emission filters for imaging with the 641 nm laser were changed slightly (680/60 bandpass filter, Omega, 655 longpass, Chroma). The light was eventually focused by the second lens in the 4f optical pathway onto a Si electron multiplying charged-coupled device camera for data and image acquisition (iXon897, Andor).

#### Calibration of silicone oil objective and comparison with standard oil objective

To calibrate the correction collar of the silicone OIO, a dilute concentration of 200 nm poly(styrene) fluorescent beads (Thermo Scientific, T7280) were immobilized in 5% (weight/volume) agarose (Invitrogen, 16520050) on a flat 200 um thick quartz coverslip (same substrate used to fabricate nanopillars). The beads were imaged in our microscope without the DHPSF mask inserted. The bead images were collected at an exposure of 50 ms and an EM gain of 200 on our microscope at ~1W/cm^2^. We fit each bead image to a 2D Gaussian with least squares regression using Matlab and extracted the peak intensity. The correction collar was set to the adjustment yielding the highest peak intensity. Although this was at the end of the adjustment range, the results were good. For comparison between objective performances, beads were immobilized in 5% agarose on the 200 um thick quartz coverslip and a 160 um thick glass substrate.

#### 2D SR data acquisition and image reconstruction

FBP17-labeled U-2 OS cells were imaged on flat glass and quartz coverslips using the standard and silicone OIOs, respectively. For both imaging configurations, the exposure time was 50 ms and the EM gain was 200. Both samples were imaged in an oxygen scavenger reductant blinking buffer to allow emitters to be confined to a long lived dark state in order to ensure sparsity. The buffer consists of 100 mM tri(hydroxymethyl)aminomethane-HCl (Thermo Fisher), 10% (weight/volume) glucose (BD Difco), 2 μL/mL catalase, 560 μg/mL glucose oxidase, and 10 mM of cysteamine (all Sigma Aldrich). The focus was set close to the coverslip near the thin edge of a cell whose membrane was spreading on the substrate in order to reduce background from out of focus emitters. A DL image at lower intensity (~1W/cm^2^) was acquired before data acquisition. After increasing the laser intensity to ~1.8 kW/cm^2^, emitters were shelved to the dark state for roughly 30 seconds. After this time period, blinking single molecules that were not overlapping were observed and data acquisition began. Roughly 40,000 frames were acquired for one cellular sample.

The data was processed in ThunderSTORM, a free ImageJ plugin. The emitters were coarsely detected with a standard maximum intensity approach. Each emitter was fit to a 2D Gaussian. The precision was calculated using Mortensen’s equation^40^. Any emitters with poor precision (> 20 nm) and whose sigma from the Gaussian fit was poor (> 200 nm) were removed. The localizations were binned in 2D histograms with a bin width of 32 nm for visualization. The widths of the tubules in the reconstructions were calculated using approaches previously described^44^.

#### 3D SR data acquisition and image reconstruction

Prior to 3D SR imaging of the samples, 200 nm poly(styrene) fluorescent beads immobilized on the surface of a flat 200 um thick quartz coverslip with 1% (weight/volume) polyvinyl alcohol (PVA) were imaged with our DHPSF microscope with the phase mask installed in the Fourier plane. The beads were imaged over an axial range of 2 um in step sizes of 50 nm at 50 ms exposure time and an EM gain of 200. We used our fine piezo XYZ stage and custom Matlab code to move the stage. This Z-scan calibration yields a curve relating lobe angle to Z position that is later used to extract Z position of emitters. Samples were incubated with dilute 200 nm poly(styrene) fluorescent beads for 8 minutes prior to imaging. The bead solution was removed and the sample was washed three times. These beads typically stick to the coverslip and provide fiducials for drift correction in post processing. The samples were then incubated in a modified blinking buffer solution consisting of 100 mM Tris-HCL, 10% glucose, 2 μL/mL catalase, 560 μg/mL glucose oxidase, and 40 mM of cysteamine and imaged. Nanopillar regions were first found by illuminating the sample with white light without the phase mask installed. In this configuration, the reference markers and arrays of pillars were clearly visible. Then, we fluorescently imaged the sample at low intensity until well-labeled samples (either nanopillars or cells on nanopillars) were visible with a fiducial in the field of view. A DL image was first taken at ~ 1W/cm^2^. Then the focus was set at the coverslip using a fiducial on the cover slip as a reference. From that point, we moved the focus 500 nm upward using the XYZ piezo stage. We inserted the phase mask, and set the laser intensity between 2.86 kW/cm^2^ and 15.9 kW/cm^2^. After 30 seconds of shelving, we acquired data of blinking DHPSFs. The exposure time was 35 ms and the EM gain was 200. We acquired approximately 70,000-100,000 frames of data.

The data was processed by fitting the emitters to a double-Gaussian function using easy-dhpsf^57^, a freely-available software in Matlab designed for localizing DHPSF emitters. The calibration curve described above was used to extract the Z positions. The precision was calculated from the detected photons using a formula calibrated for our microscope using previously described approaches^58^. After processing, poorly localized emitters (XY precision > 30 nm, Z precision > 40 nm, and lobe distance > 8 pixels) were removed. The localized single-molecule positions were rendered with the Vutara SRX program (Bruker), a software package designed for 3D SR visualization. Localizations were merged to correct for over counting (see description below). In the reconstruction, each localization was blurred by a Gaussian with a sigma of 50 nm in X, Y, and Z to reflect the localization precision in the measurements. The Z position was encoded by the color.

#### Correction of overcounting

Overcounting was corrected by first isolating clusters where the localizations in the clusters were temporally adjacent to one another. These clusters were identified by encoding the temporal information in each localization by color. Then, clusters were extracted if all the localizations in the clusters were encoded in a similar color. As the localizations in these clusters are neighboring to each other both temporally and spatially, these clusters (termed pseudo-clusters) are indicative of emitters being in consecutive frames or blinking on and off over the data acquisition. The mean off frames (number of frames between two localizations adjacent temporally) for all the clusters was 8.9. To correct for overcounting as exhibited by these pseudo-clusters, we applied a spatial and temporal threshold. This threshold ensured that any molecules within a certain spatial radius and temporal distance are merged into one molecule. We varied both the temporal and spatial threshold and observed the effect on the percentage of molecules that were merged for the surface labeled nanopillar reconstructions. After 20 off frames, the percentage of merged molecules did not change significantly, so we set the temporal threshold for all our reconstructions at 20 frames. Setting the XY distance at 50 nm and Z distance at 100 nm, we observed that the majority of the pseudo-clusters disappeared and were merged. Setting the spatial thresholds to larger values degraded the image quality as localization density throughout the reconstruction decreased. Spatial thresholds that were set lower resulted in the appearance of the pseudo-clusters. Therefore, we set the spatial thresholds to a XY distance of 50 nm and a Z distance of 100 nm for all our 3D SR reconstructions.

### Quantification and analysis

#### Diameter and curvature measurements

Using ImageJ, diameters at the center of the nanopillars were extracted from SEM images by measuring the distance of a cross section through the pillar. Only one side of the elliptical pillars was imaged for these measurements. For the surface-labeled nanopillars, individual pillars were isolated in Vutara SRX. The center of the pillar was found, and a 250 nm projection of the localizations onto the XY plane was computed. These projections were fit to an ellipse using least squares regression. The axes of the ellipse fit were used as estimations for pillar diameter. The axis of the fit that was on the same side as that for the SEM image was extracted to compare to the SEM measurements. Note, not all surface-labeled nanopillar reconstructions were analyzed. Occasionally, the fitting of the ellipse failed as there were not a sufficient number of localizations in the projection to obtain a good fit. These pillars from the surface-labeled reconstructions and SEM images were excluded in the analysis. The membrane diameter from the 3D SR reconstructions was measured with the same protocol described above. The diameters from axes of the fit of the ellipse for the membrane and surface-labeled reconstructions were averaged. Then, these averaged diameters were compared with each other yielding the results in Figure 3F.

To extract curvature from the SEM images, cross sections were measured from the bottom to the top of the nanopillars. These cross sections provided the diameters of the pillars, and the reciprocal of the radius was calculated to extract (1/R) curvature along the pillars. To calculate the curvature from the 3D SR reconstructions, points along individual nanopillars spaced evenly apart were found. These points served as the center for a 250 nm projection of the localization onto the XY plane. The projection was fit to an ellipse to extract the diameters and hence the curvature. For the surface-labeled nanopillars reconstructions, only the diameter found at the same side as the SEM image was used to calculate curvature. For the membrane reconstructions, the diameters of the fit from both axes were averaged before curvature was calculated.

#### Density calculations along nanopillars

To calculate the density of molecules along the nanopillars in our 3D SR reconstructions, we first determined the axial position of the coverslip. The coverslip was clearly apparent in the surface-labeled reconstructions. The bottom section of the membrane, the plaques, and the actin fibers were used as reference axial position of the coverslip for the cellular reconstructions. The reconstruction was then cropped from the coverslip to 1000 nm away from the coverslip axially. Then, isolated nanopillars were also cropped. These pillars were exported to custom written python code where the localizations on the pillars were projected onto the Z axis and binned in 50 nm bin widths.

#### Nanopillar simulations

To simulate the nanopillars (Figure 4D), we treated a tapering cylinder as a truncated hollow cone. The cylinder was capped with a hemi ellipsoid with a radius of 80 nm in XY and 100 nm axially. The total height of the pillar was sampled from a Gaussian distribution where the mean was 884 nm and the standard deviation was set at 71 nm, approximating the distribution of heights extracted from the SEM images of the nanopillars. The bottom diameter was set at 280 nm and the top diameter was set at 160 nm. The bottom and top diameters were chosen based on averaged values of the bottom and top diameters extracted from the top down SEM images as shown in Figure 3A (bottom right). The probability that a localization is found along the pillar was determined by the surface area of the pillar at that region. 700 localizations, roughly the average number of localizations for each nanopillar, were scattered randomly along the pillar. A Gaussian kick was added to the position of each localization laterally and axially to reflect the localization precision in our experimental measurements. The Gaussian kick distribution used a lateral sigma of 12 nm and an axial sigma of 20 nm (median experimental precisions in XY and Z). 16 total nanopillars were simulated (equivalent to the number of nanopillars experimentally analyzed in density calculations). To calculate the density of molecules along the pillar, the same analysis described in the preceding section was followed for the simulated pillars.

#### Gaussian curvature

3D localizations of isolated nanopillars from surface-labeled and membrane-labeled reconstructions were imported to the freely available package MeshLab^55^. The screened Poisson surface reconstruction^54^ was used to create a surface mesh of the imported data (see Figure S9A for a table of parameters for generating the surface). The surface was then exported as a .STL file. This file was imported to Matlab, and the Gaussian curvature was extracted along the pillar using an approach previously described^45^ using a mathematical formulation for Gaussian curvature on unstructured triangulated surfaces^56^. The Gaussian curvature along the pillar was projected into Z slices equidistant from one another. The average Gaussian curvature over all pillars was then calculated. The previous analysis was additionally used to calculate the Gaussian curvature along the simulated nanopillars.

#### Software

SEM height and diameter measurements were acquired using ImageJ. ImageJ was also used to process DL images and the 2D SR reconstructions. 3D SR reconstructions were rendered using Vutara SRX. Emitters were localized for 2D SR reconstructions using ThunderSTORM, an ImageJ plugin. Emitters were localized for 3D SR reconstructions using easy-dhpsf in Matlab. Fitting immobilized beads for objective comparison and correction collar adjustment was achieved using custom written Matlab scripts. Custom written python scripts were used for diameter and curvature measurements from SR reconstructions, for simulating the nanopillars, and for the density calculations. MeshLab was used to produce the surface meshes from the 3D localization data. Matlab was used to extract the Gaussian curvature along the surfaces.

## Supporting Information

- Correction collar calibration details, further validation of 2D and 3D SR image quality, additional 3D SR reconstructions of labeled structures, fluorescence labeling controls, localization precision distributions, figures describing methods to quantify diameters and curvature, surface meshes extracted from 3D localizations of surface-labeled reconstructions, and Gaussian curvature analysis of surface-labeled and simulated nanopillars

## Author contributions

A.R.R, W.E.M, and B.C designed experiments. A.R.R. performed all microscopy experiments and analyzed data. A.R.R. and W.Z. optimized and performed fluorescent labeling. W.Z. performed transfection and cell culture procedures. Z.J and C.T.T fabricated the nanopillars and imaged them with SEM. A.R.R. and W.E.M. wrote the initial draft. All authors edited the manuscript. W.E.M and B.C. provided conceptualization and supervision.

## Notes

W.E.M. is on the advisory board of Double-Helix Optics.

## Acknowledgments

We thank Pietro De Camilli for gifting the vector for the eGFP-FBP17 protein fusion (pEGFP-C1-FBP17). This work was supported by the National Institute of General Medical Sciences Grants R35GM118067 (to W.E.M.), R01GM128142 and R35GM141598 (to B.C.). BC grant, any fellowships ADD SNF GRANT FOR FAB

