## Supporting Information for "Exploring cell surface-nanopillar interactions with 3D super-resolution microscopy"

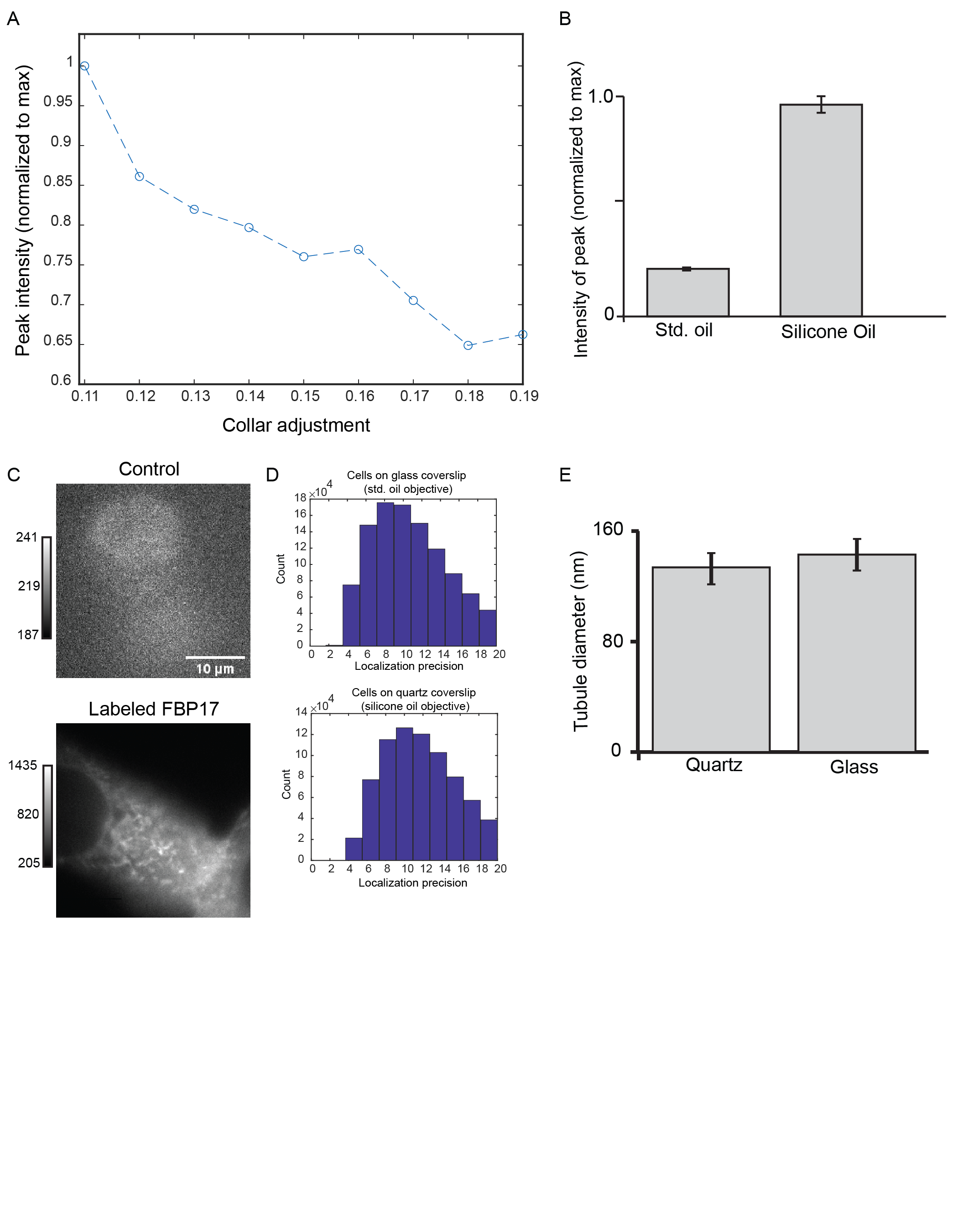

**Supporting Figure 1:** (A) Calibration of the correction collar for the silicone oil immersion objective (OIO) for a representative 200 nm fluorescent poly(styrene) bead immobilized in 5% agarose on a 200 um thick quartz substrate. Collar adjustment indicates the position that objective should be set for glass substrates at a specific thickness (ranging from 110 microns to 190 microns reflecting the collar adjustment values of 0.11 to 0.19). However, here collar must be adjusted as well as possible for a 200 um thick quartz substrate. Collar adjustment of 0.11 (maximum of curve) was used in all subsequent experiments. (B). Bar graph showing the peak intensity of beads imaged in 5% agarose on a 200 um thick quartz coverslip with the standard (std.) and silicone OIO. Error bars are ± SEM and the number of beads imaged was 30. Higher peak intensity indicates improved performance of silicone OIO. (C) Low nonspecific binding present when using AF647-GFP nanobody. Control image is a cell not transfected with GFP incubated with the GFP-AF647 nanobody. Labeled sample is a cell transfected with GFP and incubated with the GFP nanobody. Graybar indicates ADC count values on the detector. Comparison between control and labeled sample reveals low nonspecific binding. (D) Histograms of localization precisions of single molecules contributing to the 2D SR reconstructions of the labeled FBP17-eGFP cells shown in Figure 1C. (E) Bar graphs of tubule widths of FBP17 invaginations of cells grown on glass and quartz substrates imaged with the std. and silicone OIOs respectively. Error bars are SEM and number of tubules analyzed is 30.

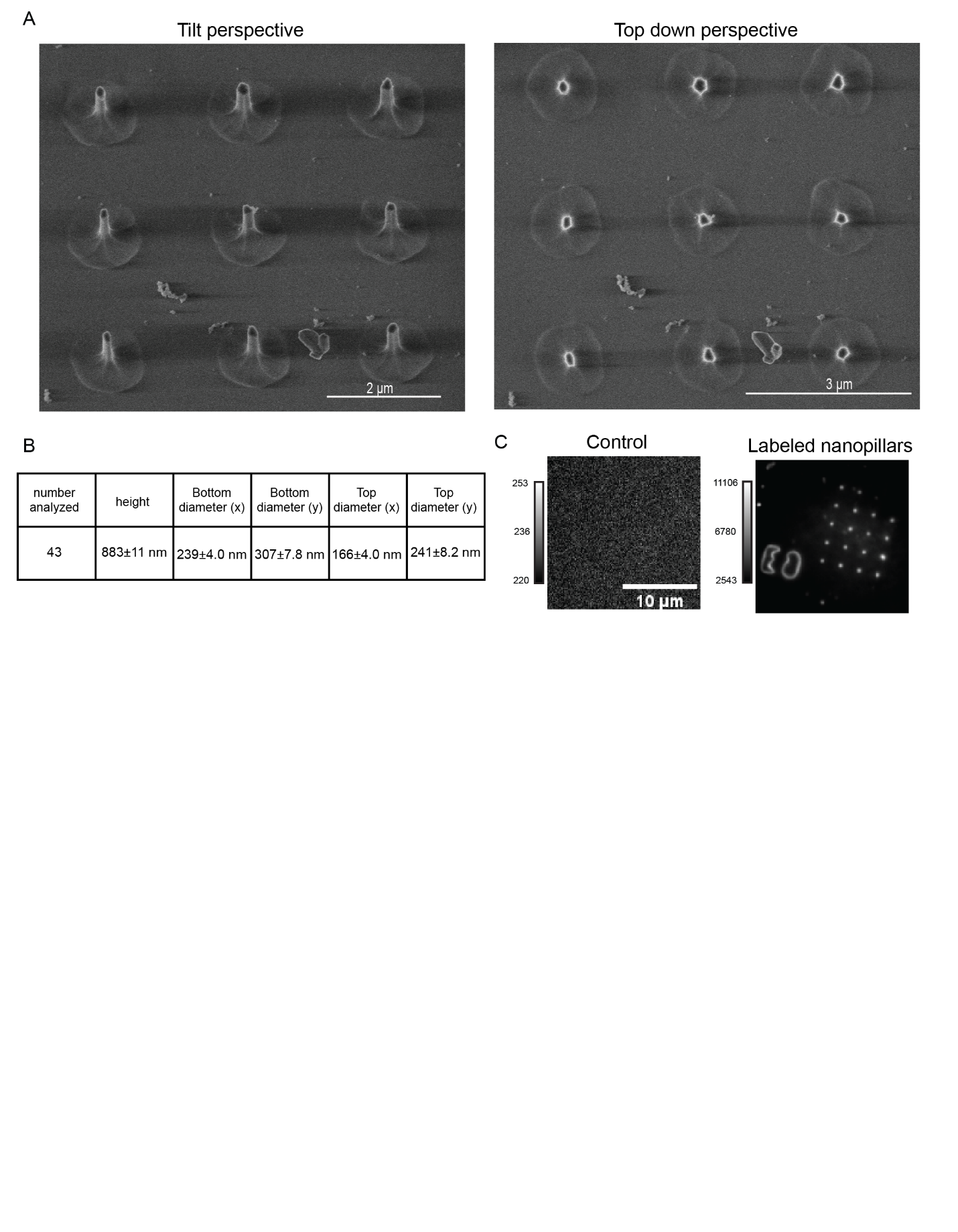

**Supporting Figure 2:** (A) SEM images of representative examples of the fabricated nanopillars. Both perspectives show the dimple near the coverslip and the elliptical shape of the nanopillars. (B) Table summarizing the measured dimensions of the nanopillars. The x direction is the direction shown in the tilted perspective of the SEM image in Figure S1A. The y direction is perpendicular to the x direction. The diameters were extracted from the top down perspective. Values are mean ± SEM. (C) Low nonspecific binding present when labeling pillars with AF647 NHS ester. Control sample depicts a nanopillar region incubated with AF647 NHS ester where pillars were not incubated with (3-Aminopropyl) triethoxysilane. The treatment with (3-Aminopropyl) triethoxysilane forms free lysine residues on the surface of the pillars that will then react with AF647 NHS ester to covalently attach the fluorophore. The labeled nanopillars were incubated with both AF647 NHS ester and (3-Aminopropyl) triethoxysilane.

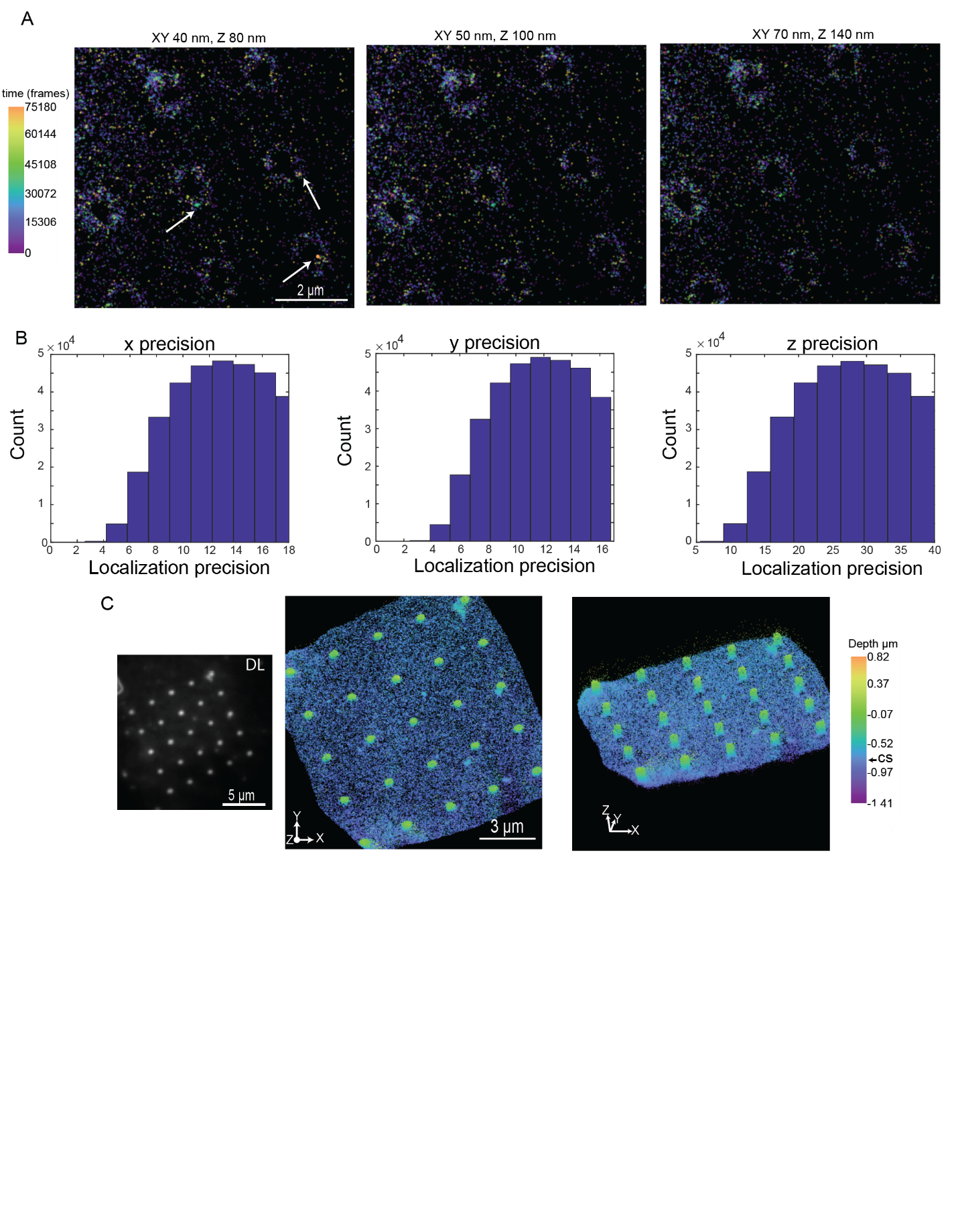

**Supporting Figure 3:** (A) Merging single molecules by varying the spatial threshold in 3D SR reconstructions of surface labeled nanopillars. The spatial threshold (shown at the top) was varied for a 3D SR reconstruction of surface labeled nanopillars. The temporal threshold was set at 20 frames. The images depict the pillars near the coverslip and each localization is temporally encoded by color. White arrows in far left image show bright clusters where the localizations are close to one another in time. These are pseudo-clusters that are a result of localizing a single molecule more than once. Increasing spatial threshold (middle) shows a decrease in the size and frequency of the pseudo-clusters. Increasing spatial threshold further (right) appears to decrease density around pillars. Thus, the middle spatial threshold was chosen and applied to all 3D SR reconstructions. (B) Histograms showing the X, Y, and Z localization precision of the single molecules contributing to the 3D SR reconstruction shown in Figure 3B. (C) Additional 3D SR reconstruction of surface labeled nanopillars. This 3D SR reconstruction is in the same array pattern as the DL image and shows the cylindrical pillar structures.

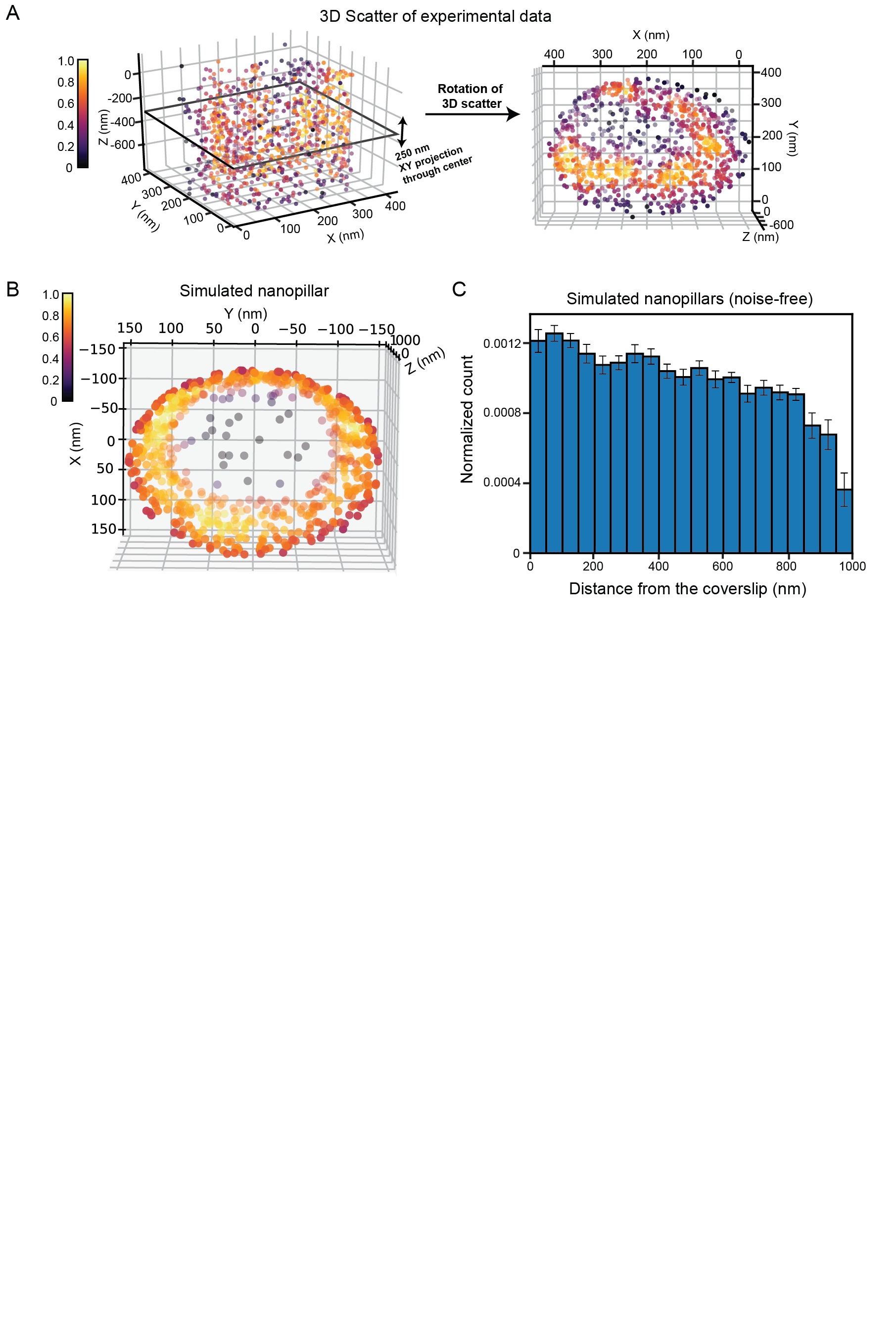

**Supporting Figure 4:** (A) 3D density plot of a nanopillar depicting extraction of the diameter measurement. The localizations are color encoded by their local density. The parallelogram at the center represents the position of 250 nm thick XY projection of the localizations. These localizations are then fit to an ellipse to extract the diameter. Rotation of the 3D scatter (right) shows the bottom of the nanopillar with the localizations clearly forming a hollow ring. Grey localizations in the center are from the top of the pillar. (B) 3D density plot of a simulated nanopillar. Pillar is shown in a bottom view with the ring clearly evident. Grey localizations at the center are localizations at the top. (C) The simulated nanopillars (n=16) were projected into 50 nm Z slices along the nanopillars and binned into a histogram. No Gaussian kicks reflecting the localization precision of measurement were added to these simulated nanopillar. The shape of the distribution of normalized counts decreases closer to the top as expected for tapering nanopillars. The shape is similar to the experimental distribution and the distribution extracted from simulated nanopillars where Gaussian kicks were added. Error bars are SEM.

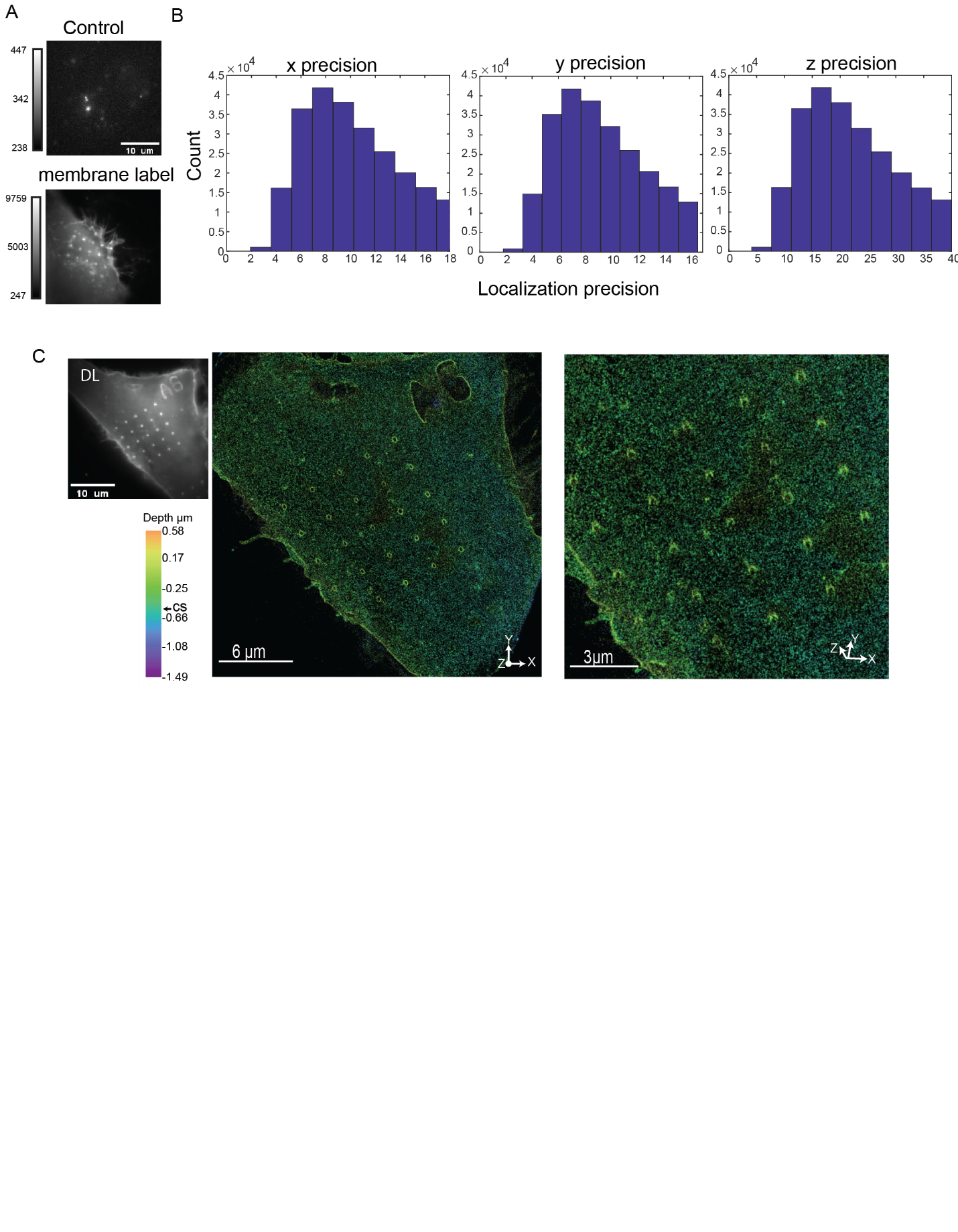

**Supporting Figure 5:** (A) Low nonspecific binding of SNAP-surface AF647. Control sample depicts a cell adhering on nanopillars incubated with SNAP-surface AF647 but not transfected with the pDisplay SNAPtag vector. The membrane label case shows a cell adhering to nanopillars incubated with SNAP-surface AF647 and transfected with the pDisplay SNAPtag vector. (B) Distributions of X, Y, and Z localization precision from the 3D SR reconstruction of the transmembrane labeled cell depicted in Figure 5A. (C) Additional 3D SR reconstruction of a transmembrane labeled cell adhering to the nanopillar. The pillar array in the reconstruction follows the same pattern in the DL image. Cylindrical pillar structures are visible in the tilted image.

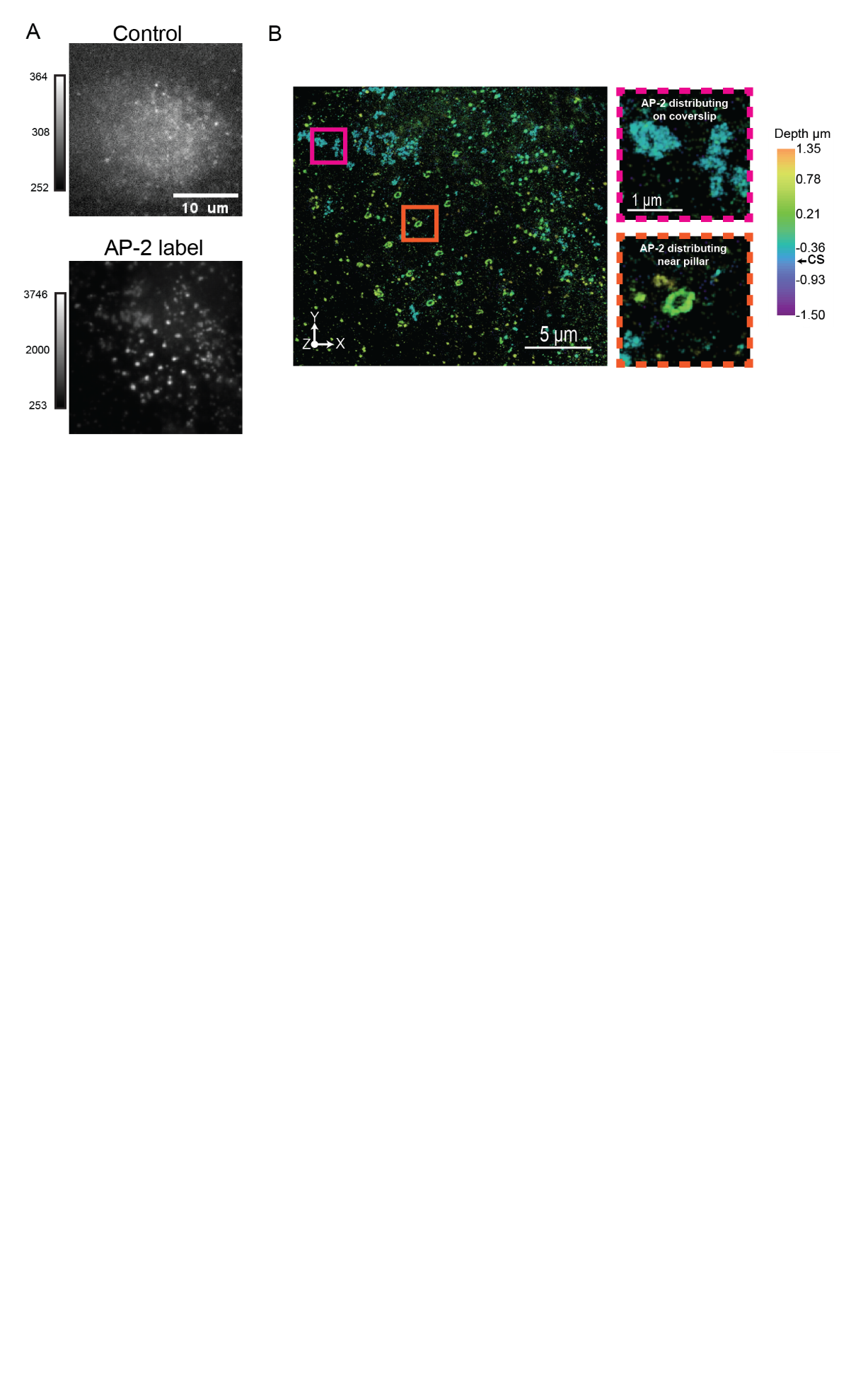

**Supporting Figure 6:**  Low nonspecific binding of the secondary antibody attached to AF647 used in AP-2 imaging. The control sample depicts a cell adhering on nanopillars but not incubated with the primary antibody during sample preparation. The AP-2 label case shows a cell adhering to the nanopillar and incubated with the primary antibody. (B) Additional 3D SR reconstruction of an AP-2 labeled cell adhering to the nanopillars. The pillar array is similar to the pattern in the DL image. Insets show AP-2 distributing on the coverslip and distributing around a nanopillar. Lack of teal color at nanopillar reveals that AP-2 does not localize near the bottom of the nanopillars.

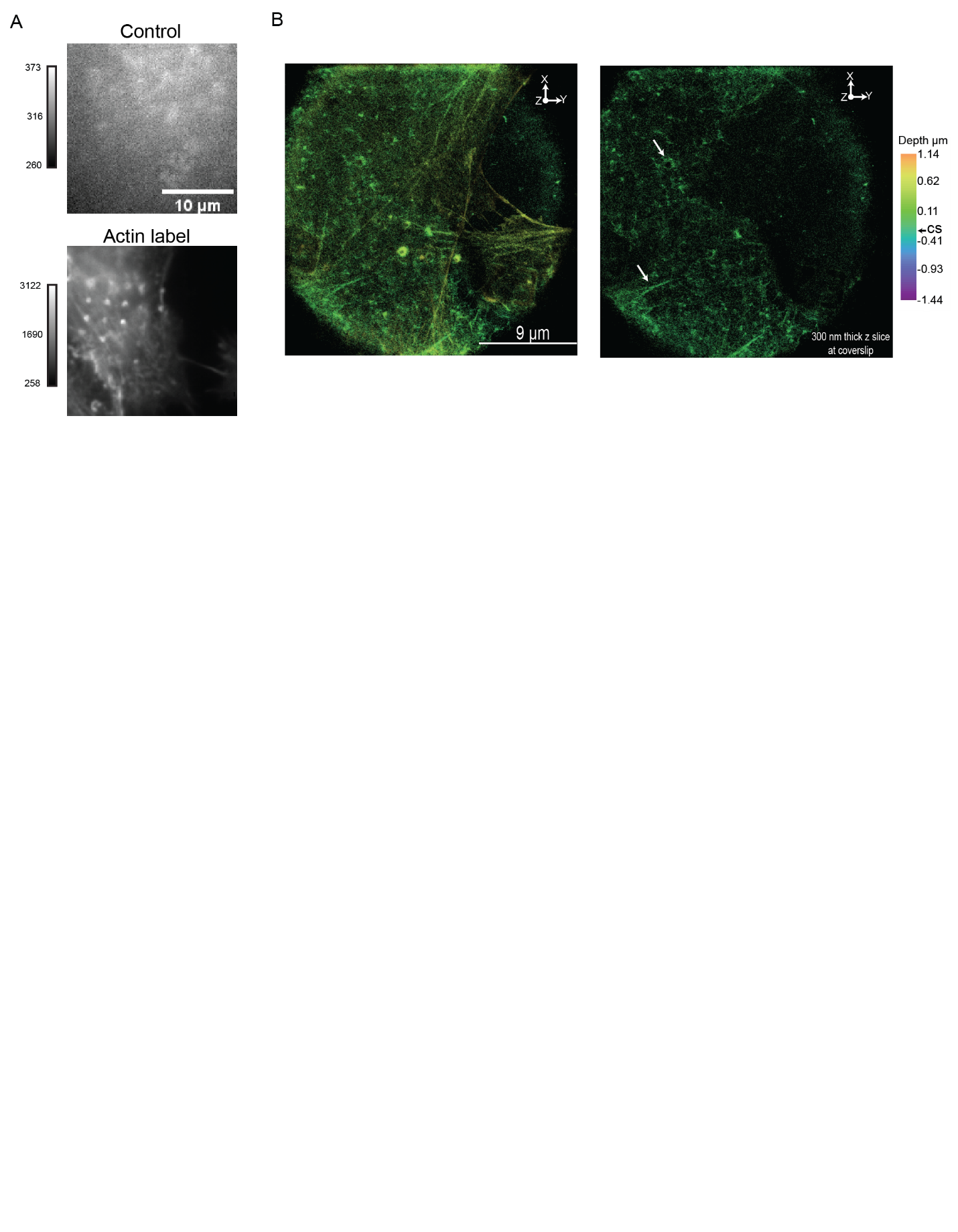

**Supporting Figure 7:**  Low nonspecific binding of phallodin-AF647. The control sample depicts a nanopillar region incubated with phallodin-AF647. The actin label case shows a cell adhering to the nanopillar incubated with the phallodin-AF647. (B) Additional 3D SR reconstruction of actin labeled cells adhering to the nanopillars. The pillar array is similar to the pattern in the DL image shown in A. The image on the right is a 300 nm Z slice close to the coverslip. The white arrows depict actin fibers and actin accumulating around the nanopillars.

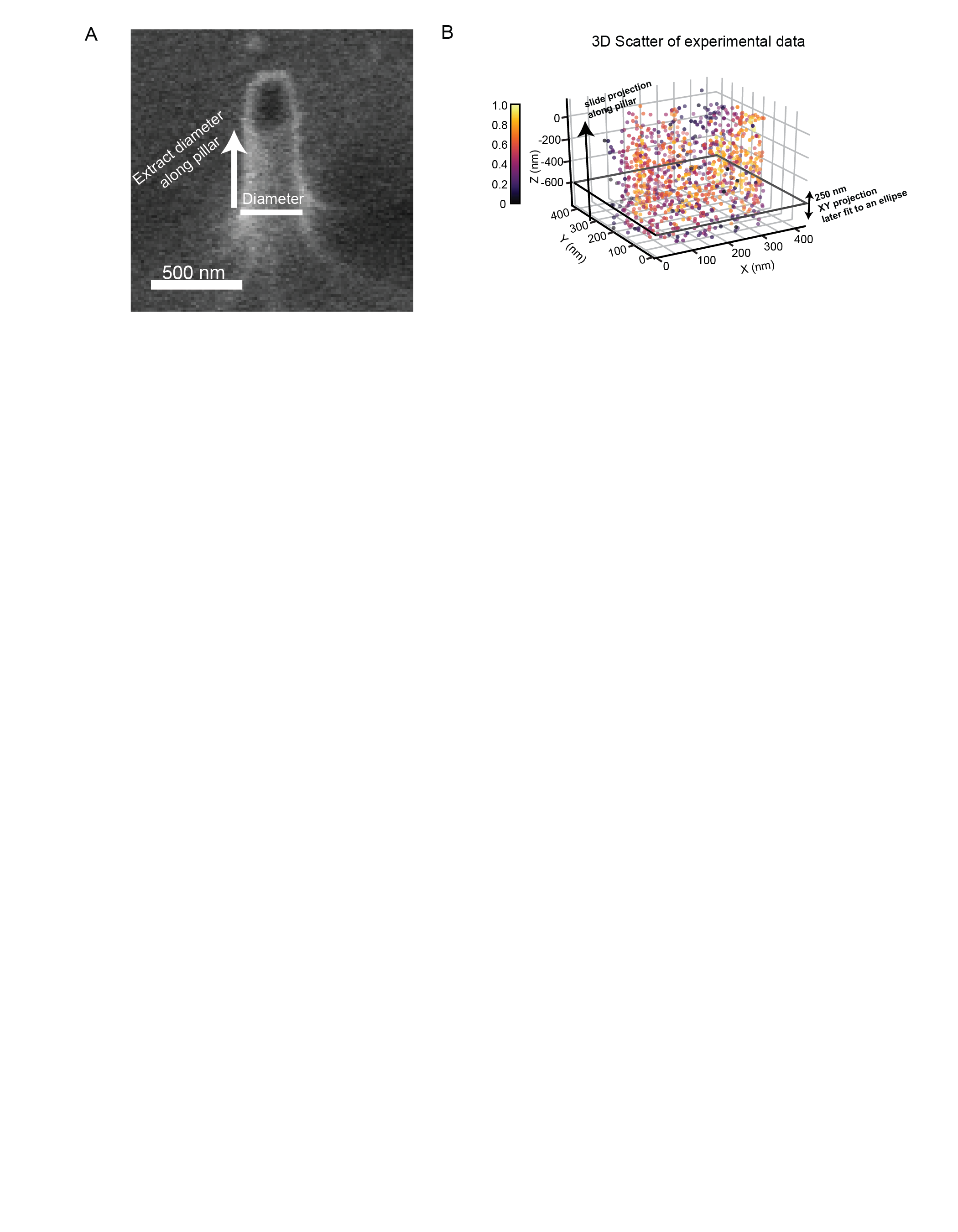

**Supporting Figure 8:** (A). Extracting curvature from SEM images. The SEM image depicts the protocol to extract the diameter along the nanopillar. A cross section is first taken at the bottom and later at specific points along the pillar. The reciprocal of the radius is then calculated to extract the 1/R curvature. (B). Extracting the diameters from the 3D SR scatter plots. A 250 nm XY projection near the bottom of the pillar is fit to an ellipse to extract the diameters of the pillars. This projection is then slid to various axial points along the pillar. From the fit, the reciprocal of the radius is calculated to find the curvature of the pillar. The 3D scatter shown is from a surface labeled nanopillar reconstruction. This method is also used to calculate the 1/R curvature from the transmembrane reconstructions.

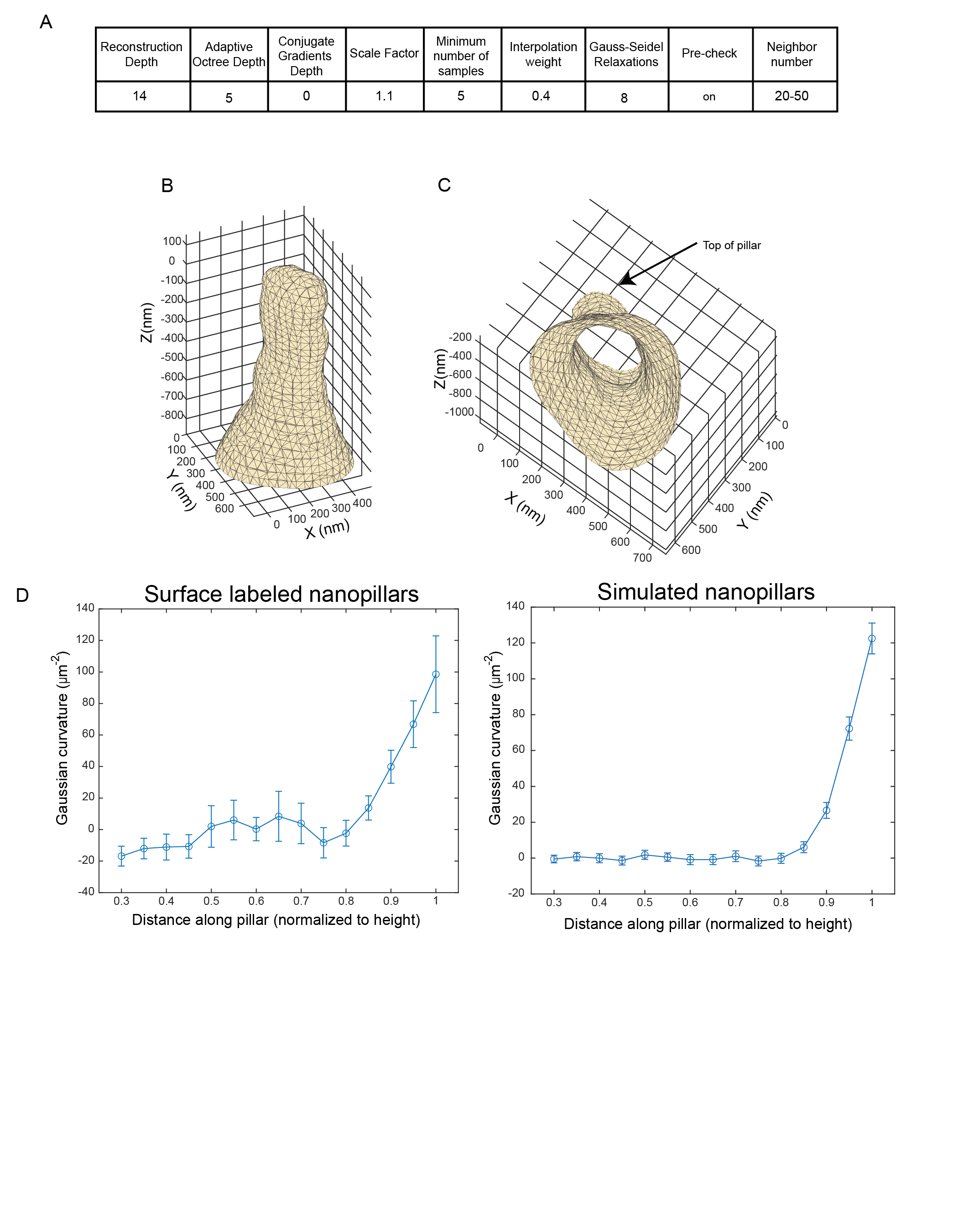

**Supporting Figure 9:** (A). Parameters used to generate surface mesh. The screened Poisson surface reconstruction algorithm in MeshLab was used to generate surface meshes from the 3D localization data of surface labeled and transmembrane labeled reconstructions. (B). Surface mesh of a pillar from the surface labeled nanopillar reconstructions. (C) Rotation of the same pillar reveals that the pillar is hollow. The top cap of the pillar has been removed to clearly showcase the hollow features of the mesh. (D) Gaussian curvature calculations with axial location along surface labeled (left) and simulated (right) nanopillars. The Gaussian curvature was calculated along (n=15) surface labeled and simulated nanopillars as described in the methods section. Error bars are SEM. We clearly observe close to zero Gaussian curvature along the shaft of the pillar and highly positive Gaussian curvature near the top.
